# Comparative genomics of the extremophile *Cryomyces antarcticus* and other psychrophilic Dothideomycetes

**DOI:** 10.1101/2024.04.29.590463

**Authors:** Sandra V. Gomez, Wily Sic, Sajeet Haridas, Kurt LaButti, Joanne Eichenberger, Navneet Kaur, Anna Lipzen, Kerrie Barry, Stephen B. Goodwin, Michael Gribskov, Igor V. Grigoriev

## Abstract

*Cryomyces antarcticus* is an endolithic fungus that inhabits rock outcrops in Antarctica. It survives extremes of cold, humidity and solar radiation in one of the least habitable environments on Earth. This fungus is unusual because it produces heavily melanized, meristematic growth and is thought to be haploid and asexual. Due to its growth in the most extreme environment, it has been suggested as an organism that could survive on Mars. However, the mechanisms it uses to achieve its extremophilic nature are not known. Over a billion years of fungal evolution has enabled representatives of this kingdom to populate almost all parts of planet Earth and to adapt to some of its most uninhabitable environments including extremes of temperature, salinity, pH, water, light, or other sources of radiation. Comparative genomics can provide clues to the processes underlying biological diversity, evolution, and adaptation. This effort has been greatly facilitated by the 1000 Fungal Genomes project and the JGI MycoCosm portal where sequenced genomes have been assembled into phylogenetic and ecological groups representing different projects, lifestyles, ecologies, and evolutionary histories. Comparative genomics within and between these groups provides insights into fungal adaptations, for example to extreme environmental conditions. Here, we analyze two *Cryomyces* genomes in the context of additional psychrophilic fungi, as well as non-psychrophilic fungi with diverse lifestyles selected from the MycoCosm database. This analysis identifies families of genes that are expanded and contracted in *Cryomyces* and other psychrophiles and may explain their extremophilic lifestyle. Higher GC contents of genes and of bases in the third positions of codons may help to stabilize DNA under extreme conditions. Numerous smaller contigs in *C. antarcticus* suggest the presence of an alternative haplotype that could indicate that the sequenced isolate is diploid or dikaryotic. These analyses provide a first step to unraveling the secrets of the extreme lifestyle of *C. antarcticus*.

## 1 Introduction

Life in extreme cold and high-altitude environments like Antarctica and the Alps may be severely restricted, but it is not entirely absent. Under these conditions, rocks are the principal habitat for microorganisms, which are known as epilithic when growing on rock surfaces or endolithic when growing inside the substrate. Endolithic organisms inhabiting rock are categorized into chasmoendoliths, which inhabit fissures and cracks, and cryptoendoliths, which inhabit porous rock substrates (Friedmann, 1982). Cryptoendoliths comprise filamentous fungi, unicellular green algae, and bacteria. Fungi growing under these conditions are known as microcolonial fungi (MCF) or rock-inhabiting fungi (RIF). Most of these organisms belong to the order Dothideomycetes but some are placed in the Eurotiomycetes or remain taxonomically undetermined (Ruibal *et al*., 2009; Selbmann *et al*., 2015).

Cryptoendolithic species in genera such as *Cryomyces* and *Friedmanniomyces* have thick, melanized cell walls, produce exopolysaccharides (extracellular polymeric substances, or EPS) outside the hyphae, and have other adaptations to harsh environments (Selbmann *et al*., 2005). Physiological adaptations such as the production of trehalose and cryoprotectant sugars, polyols, lipids/fatty acids, antifreeze proteins (Robinson, 2001, Ayukawa *et al*., 2016), and hydrolases and oxidoreductases (Duarte *et al*., 2018, Coleine et al., 2022) play an important role in the adaptation of these organisms to cold. In general, psychrophilic black fungi show increased protein expression when grown at 1 C; *Friedmanniomyces endolithicus*, one of the species included in this work, shows increased numbers of high-molecular weight proteins in comparison to mesophilic fungi (Tesei *et al*., 2012), suggesting substantial reprogramming of protein expression. These and other physiological adaptations to extreme environments should be reflected in genome modifications.

The production of proteins and enzymes underlies the physiological mechanism of cold tolerance in fungi. Observed changes occur in both gene regulation and as mutational changes to individual proteins. One of the simplest adaptations is the expansion or contraction of paralogous groups. Changes in gene number provide a rapid means to modify proteome composition, enzymatic activity, and the constituents of the membrane, all of which are important in cold/stress adaptation. However, these mechanisms are complex and not fully understood (Maggi *et al*., 2013).

Adaptation to cold environments may require the alteration of proteins such as those related to membrane transport and lipid biosynthesis to maintain membrane flexibility at low temperatures. A proteome comparison can identify proteins that are significantly altered in cold-adapted fungi. However, studies of proteome composition mostly have been limited to the comparison of only a few species at a time, limiting our ability to see general proteomic patterns related to adaptation and survival in extreme conditions. Identification of such protein and enzyme patterns in the proteomes of cold-adapted fungi can identify proteins that are responsible for physiological mechanisms of cold tolerance.

Expansion or contraction of orthologous protein families is one of the simplest ways for organisms to adapt to an extreme environment. Expansion of specific groups has been identified in many species (Su *et al*., 2016; Stajich, 2017; Paiva *et al*., 2024). Previous analysis of a draft genome of *C. antarcticus,* sequence with Ion PI, identified little that stood out (Sterflinger et al., 2014), but that analysis included comparisons with only five other species, only three of which were in the Dothideomycetes, and the assembly was very fragmented. In this work, we compare the proteome composition of 52 species in the class *Dothideomycetes*, 19 of which are extremophiles (cold, acidophilic, ethanol vapor, xerophilic, or salt tolerant). The selected species include 11 psychrophiles which are the primary targets of our analysis. The non-psychrophilic species include lichens, yeasts, parasitic species, opportunistic human pathogens, and plant pathogens from selected Dothideomycetes groups. We examine and compare the expansion and contraction of orthologous groups to test the hypothesis that families of genes associated with adaptation may be altered in these stress-tolerant Dothideomycetes.

Based on previous published work (reviewed by Gostinčar and Gunde-Cimerman, 2023), we expect to see changes in genes related to pigment production, cell wall biosynthesis, membrane transporters, production of compatible solutes (such as glycerol), fatty acid synthesis, and response to reactive oxygen species (ROS). In addition, specific groups necessary for survival in other environments, for instance genes involved in lignin degradation and carbohydrate-active enzymes (CAZymes) required for pathogenicity to plants, may be dispensable in environments where the requisite substrates are unavailable. By synthesizing results across a broad range of species, we hope to distinguish general principles of adaptation to extremely cold environments from idiosyncratic adaptation seen in only one or two species.

## 2 Materials and methods

### 2.1 Genome sequencing, assembly, and annotation

*Cryomyces antarcticus* isolate 116301 was obtained from the CBS culture collection (Utrecht, the Netherlands). The fungus was cultured on 2% potato dextrose agar plates at 4 C for more than 6 months. Mycelia were scraped off the surface of the plates and lyophilized. DNA was extracted using Qiagen’s DNeasy kit and RNA was extracted using the RNeasy kit. Its genome was sequenced using the PacBio platform. For the PacBio library, 5 μg of genomic DNA was sheared to >10 kb using Covaris g-Tubes. The sheared DNA was treated with exonuclease to remove single-stranded ends followed by end repair and ligation of blunt adapters using the SMRTbell Template Prep Kit 1.0 (Pacific Biosciences). The library was purified with AMPure PB beads. PacBio Sequencing primer was then annealed to the SMRTbell template library and sequencing polymerase was bound to them using the Sequel II Binding kit 1.0. The prepared SMRTbell template libraries were then sequenced on a Pacific Biosystems Sequel II sequencer using 8M v1 SMRT cells and Version 1.0 sequencing chemistry with 1×900 min sequencing movie run times. Filtered subread data were processed with the JGI QC pipeline to remove artifacts and organellar reads. Filtered Circular Consensus Sequence (CCS) reads were assembled with Flye version 2.7.1-b1590 (Kolmogorov *et al*., 2019) to generate an assembly and polished with the PacBio SMRTLINK (v8.0.0.80529) using the gcpp--algorithm arrow command.

For the *Cryomyces antarcticus* transcriptome, plate-based RNA sample prep was performed using the PerkinElmer Sciclone NGS robotic liquid handling system and Illumina’s TruSeq Stranded mRNA HT sample prep kit, using poly-A selection of mRNA following the protocol outlined by Illumina in their user guide (https://support.illumina.com/sequencing/sequencing_kits/truseq-stranded-mrna.html) and with the following conditions: total RNA starting material was 1 μg per sample and 8 cycles of PCR were used for library amplification. The prepared library was then quantified using KAPA Biosystem’s next-generation sequencing library qPCR kit (Roche) and run on a Roche LightCycler 480 real-time PCR instrument. The quantified library was then multiplexed with other libraries, and the pool of libraries was prepared for sequencing on the Illumina NovaSeq 6000 sequencing platform using NovaSeq XP v1 reagent kits, S4 flow cell, following a 2×150 indexed run recipe. Raw RNAseq reads were filtered and trimmed using the BBDuk kmer matching (kmer=25) and phred trimming method set at Q6. Following trimming, reads under the length threshold were removed (minimum length of 25 bases or 1/3 of the original read length - whichever is longer). Filtered reads were used as input for *de novo* assembly of RNA contigs using Trinity version 2.8.5 with parameters--normalize_reads and --jaccard_clip.

The genome was annotated using the JGI Annotation pipeline and made available at the JGI MycoCosm genome portal (http://mycocosm.jgi.doe.gov/Cryan3) along with tools for interactive comparative analysis (Grigoriev et al., 2014). Both the assembly and annotation have been deposited to GenBank under accession (TO BE PROVIDED UPON PUBLICATION).

### 2.2 Obtention of genome data

The proteomes of 52 fungal species were obtained from the JGI MycoCosm portal (https://mycocosm.jgi.doe.gov/mycocosm/; Grigoriev *et al*., 2014) (Table 1). The majority of the species belong to the subclass Dothideomycetidae within the Dothideomycetes. Members of this subclass were chosen for their close phylogenetic relationship to *Cryomyces antarcticus* and *Cryomyces minteri*, as indicated by the phylogenetic tree available in MycoCosm (https://mycocosm.jgi.doe.gov/mycocosm/species-tree/tree;j7jS2o?organism=dothideomycetes).

**Table 1.**
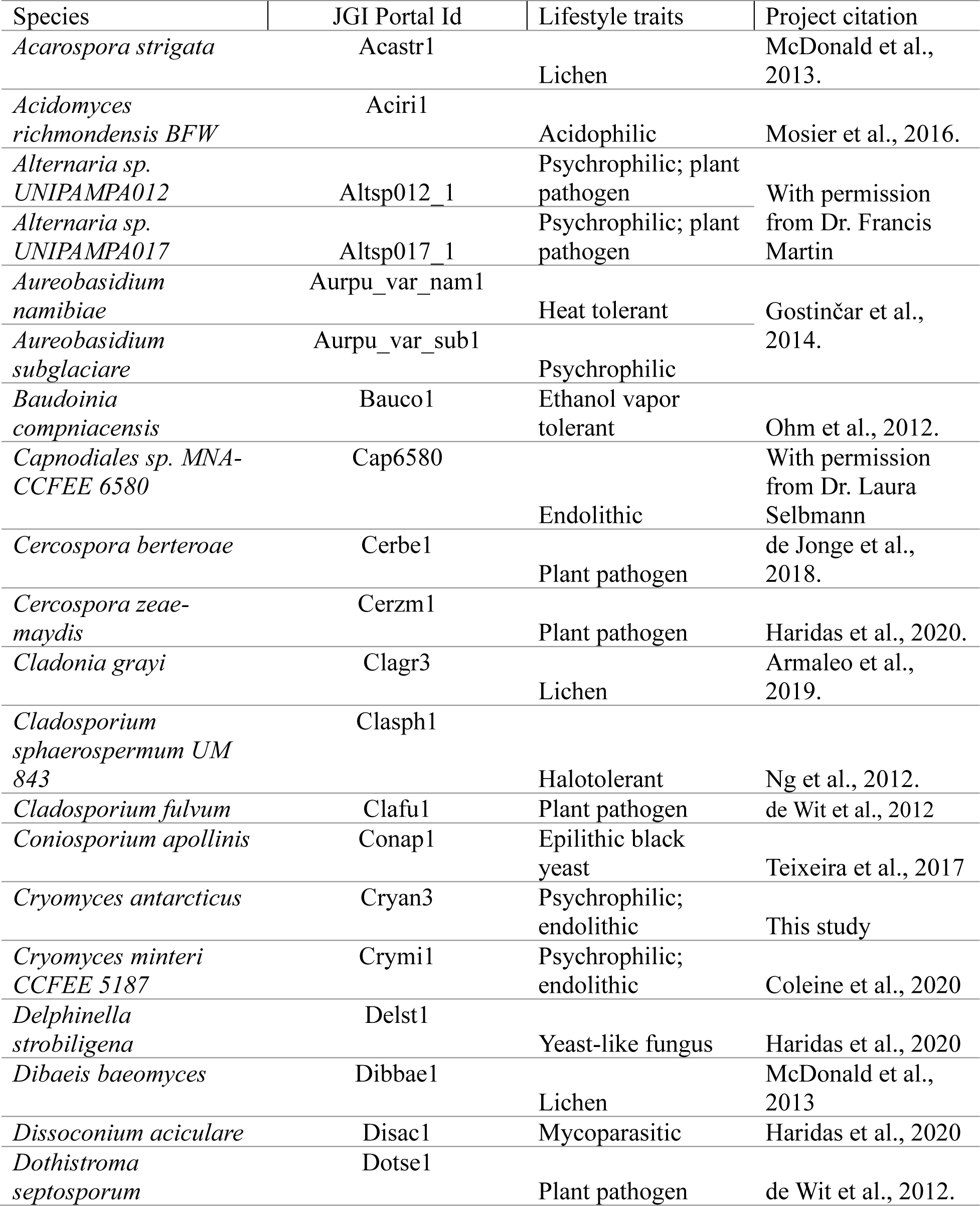
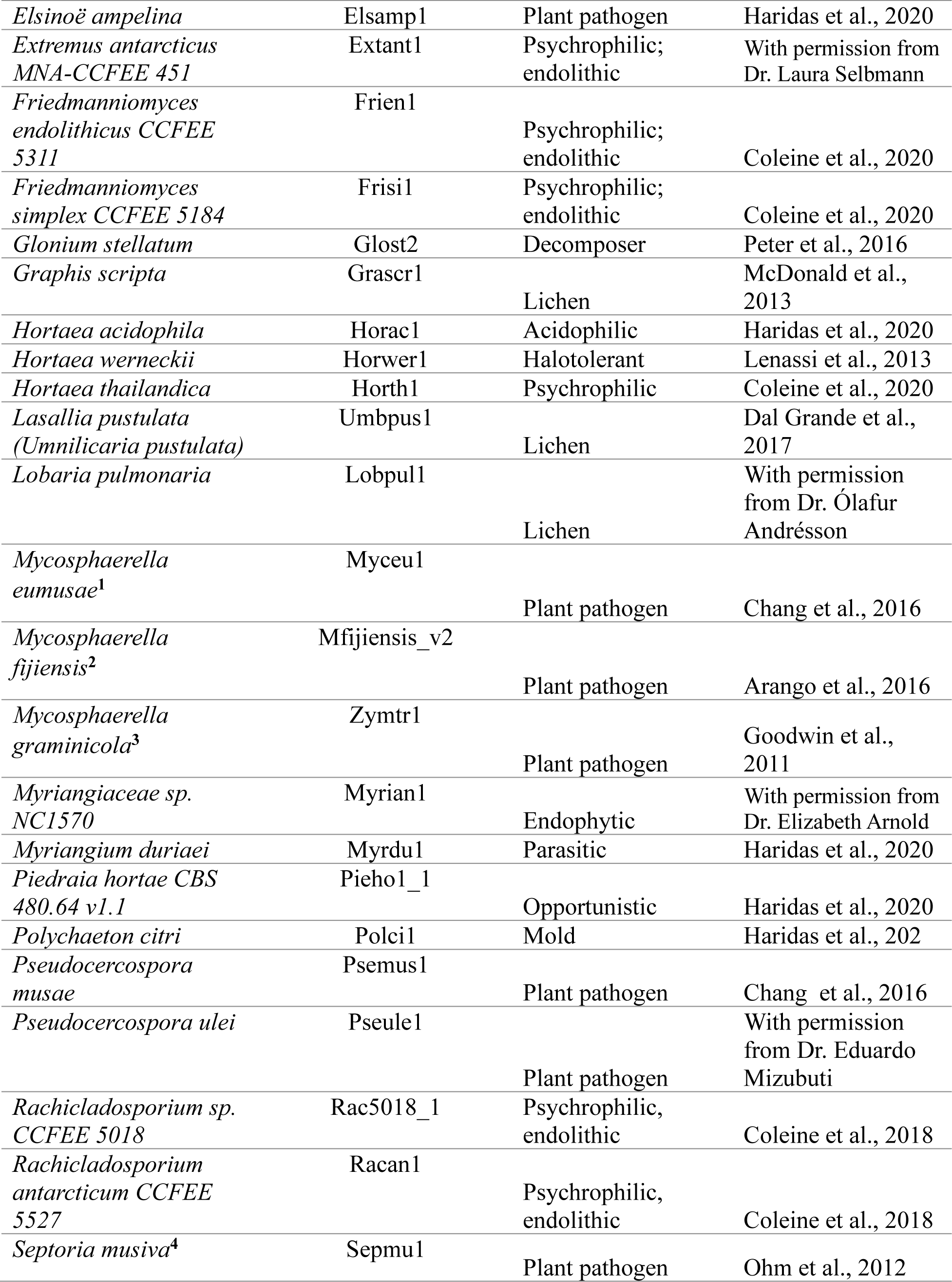
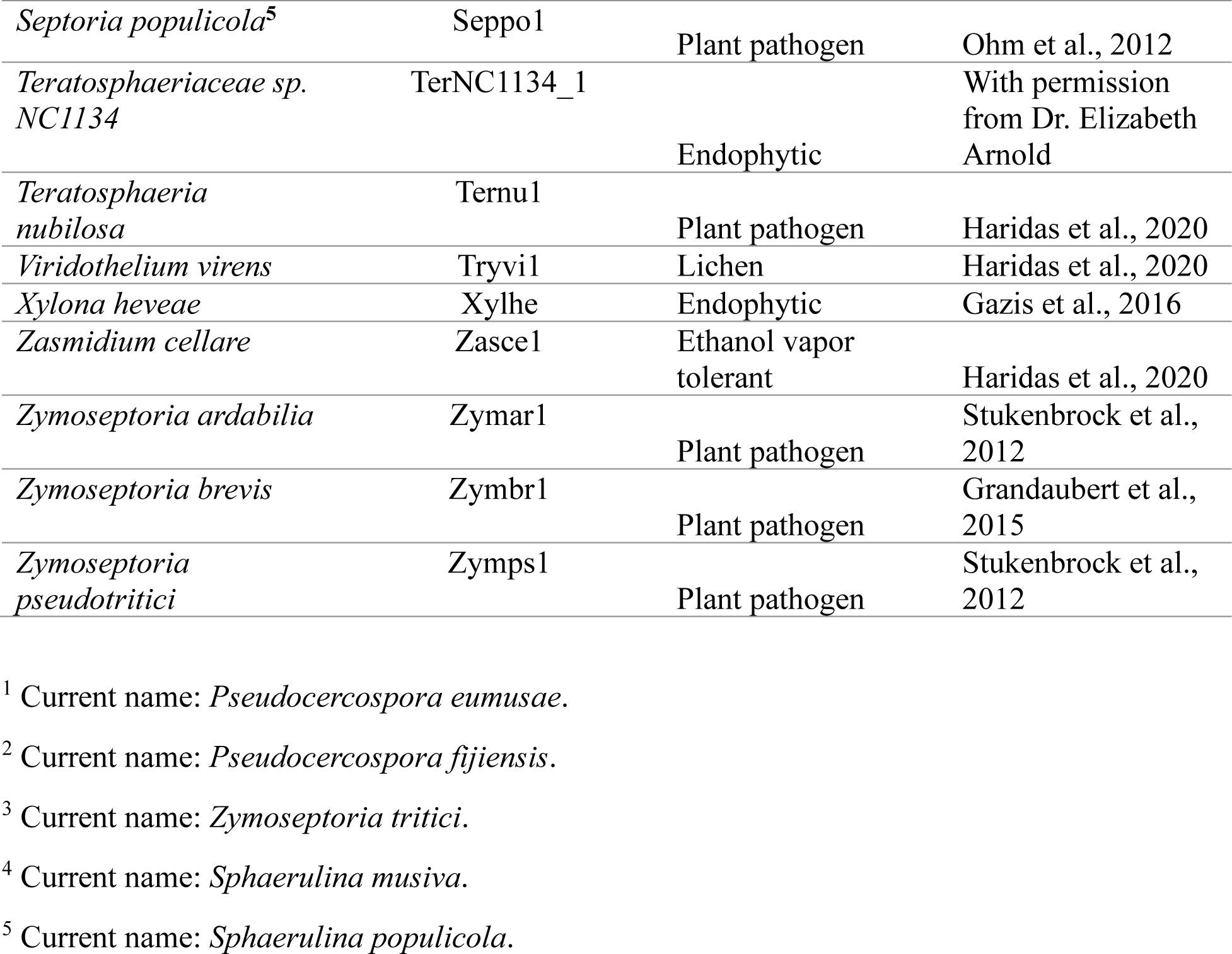
Summary information about the Dothideomycetes fungal species included in the comparative genomic analyses.

The group of psychrophilic fungi (https://mycocosm.jgi.doe.gov/Psychrophilic_fungi) consists of 11 species, including our target species *C*. *antarcticus* and *C. minteri*, as well as the species *Aureobasidium subglaciare, Extremus antarcticus, Friedmanniomyces simplex, Hortaea thailandica, Rachicladosporium* sp., *Rachicladosporium antarcticum,* and two isolates of the genus *Alternaria* from the order Pleosporales, *Alternaria* sp. UNIPAMPA012 and *Alternaria* sp. UNIPAMPA017. Both these species were isolated from healthy leaves of Antarctic hair grass.

Among the selected species, we include representatives of different lifestyles: endophytic, endolithic, molds, yeasts, and opportunistic human pathogens, parasites, and plant pathogens (Table 1). The non-psychrophilic group contains fungal pathogens of a diverse range of hosts including the sugar beet pathogen *Cercospora berteroae*, the corn pathogen *Cercospora zeae-maydis*, the tomato and grape pathogens *Cladosporium fulvum* and *Elsinoë ampelina*, the banana pathogens *Pseudocercospora* (formerly *Mycosphaerella*) *eumusae*, *P*. (*Mycosphaerella*) *fijiensis*, and *P. musae*, the wheat pathogen *Zymoseptoria tritici* (formerly *Mycosphaerella graminicola*) and its close relatives *Z. ardabiliae*, *Z. brevis*, and *Z. pseudotritici* (isolated from wild grass species). We also include the forest pathogens *Dothistroma septosporum, Pseudocercospora ulei, Septoria musiva* (now *Sphaerulina musiva*; teleomorph: *Mycosphaerella populorum*)*, Septoria populicola,* and *Teratosphaeria nubilosa*. In addition, other extremophiles are included in the analysis to compare the survival strategies under different extreme environmental conditions: the acidophilic fungi *Acidomyces richmondensis* and *Hortaea acidophila;* the heat-tolerant *Aureobasidium namibiae;* the ethanol vapor and heat-tolerant species *Baudoinia compniacensis* and *Zasmidium cellare*; and the salt-tolerant species *Hortaea werneckii* and *Cladosporium sphaerospermum*. Furthermore, six species of lichens (*Acarospora strigata*, *Cladonia grayi*, *Dibaeis baeomyces, Graphis scripta, Umbilicaria pustulata* and *Lobaria pulmonaria*) from the class Lecanoromycetes, and one endophytic species (*Xylona heveae*) from the Xylonomycetes, were included as outgroups.

### 2.3 GC contents of the genomes, coding sequences and third-codon position (GC3)

GC content was analyzed using the stats.sh function of bbtools (Bushnell, 2014) for determining the overall GC contents of the whole-genome sequences, and the EMBOSS application “cusp” (Rice et al., 2000) for calculating the GC content of protein-coding genes and the third-codon position (GC3). GC content of genomic contigs and scaffolds of *C. antarcticus* and *C. minteri* were calculated using countgc.sh of the bbtools package (Bushnell, 2014). Plots representing the GC distribution were created using ggplot2 (Wickham, 2016) in R.

### 2.4 Genome-wide analysis of repeat-induced point mutation (RIP)

Genome assemblies of species included in this study were used to conduct RIP analyses using the RIPper software (https://github.com/TheRIPper-Fungi/TheRIPper/) (van Wyk *et al*., 2019). This program calculates the RIP indices and the average GC content of whole-genome sequence information using a sliding-window approach. Large RIP-affected genomic regions (LRARs) and the index values associated with them are also determined. We used the default parameters for genome-wide RIP index calculations consisting of 1000-bp windows with a 500-bp step size. Stringent parameters were used (product index value > 1.15; substrate index value < 0.75; RIP composite index > 0) to identify genomic regions affected by RIP. RIPper considers a window as RIP positive if all three indices indicate RIP activity. Only regions of more than 4000 consecutive bp that are affected by RIP are considered as LRARs (van Wyk *et al*., 2019). The total estimated genome-wide RIP percentage and the composite index for LRARs are shown along with a rooted species tree that was constructed using OrthoFinder v. 2.5.5 (Emms and Kelly, 2019) and the proteome files of each species. The species phylogenetic tree along with the RIP analysis results were visualized using the Interactive Tree of Life (iTOL v. 5.0) (Letunic and Bork, 2021).

### 2.5 Identification of orthogroups

OrthoFinder v. 2.5.5 (Emms and Kelly, 2019) was used to identify orthologous gene families using DIAMOND (Buchfink, Reuter and Drost, 2021) and the MCL graph clustering (Enright, Van Dongen and Ouzounis, 2002) for sequence comparison and STAG (Emms and Kelly, 2018) as the default species tree method. For each of the 25,761 orthogroups, gene counts per species were normalized by converting to standard normal deviates using the 50% trimmed mean and standard deviation of each orthogroup. Expanded and contracted orthogroups were identified by comparing the normalized scores for the 11 psychrophiles and the 41 non-psychrophiles using Welch’s t-test statistic,

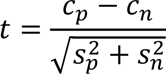

where c is the mean normalized count, s is the standard deviation, and n is the number of sequences. Subscripts *p* and *n* indicate psychrophiles and non-psychrophiles, respectively. The 50 most expanded and 50 most contracted groups were selected for further analysis.

### 2.6 Correlational matrix and heatmap

The R built-in stats function “cor” (R Core Team, 2013) and “reshape2” (Wickham, 2007) package were used to build a correlation matrix using the normalized counts. We used the package “factoextra” (Kassambara and Mundt, 2020) in R to conduct a principal component analysis (PCA) using the correlational matrix to determine clusters of species explained by the orthogroups. The packages “ggplot2” and “ggcorrplot” (R Core Team, 2013) were used to visualize the associations in the matrix.

The R package *pheatmap* (v1.0.12) (Kolde, 2015) was used to construct a heatmap of the selected expanded and contracted orthogroups. Correlation was used as the distance measure for both orthogroups and species, and groups were clustered using Ward’s method (Ward Jr., 1963).

### 2.7 Functional annotation of orthogroups

Orthogroup annotations were obtained by Blast comparison to the UniRef50 database (The UniProt Consortium, 2023) with a maximum E-value of 10^−10^ and minimum of 40% identity. To identify domain annotations and GO terms associated with orthogroups, we used Interproscan (v 5.66-98.0) and InterPro (v 98.0) (Paysan-Lafosse *et al*., 2023). CAZymes were annotated using dbCAN2 (Yin *et al*., 2012).

## 3 Results

### 3.1 Genome features of psychrophilic fungi

Genome sizes of the psychrophilic fungi range from 23.88 to 51.66 Mbp, with *C. antarcticus* being the largest in the list (Figure 1). The number of encoded proteins varies from 8,778 to 18,892, with 9,404 in *C. antarcticus* (Figure 2). A VISTA dot plot of the *C. antarcticus* contigs revealed numerous diagonal lines (Figure 3; Suppl. Figure 1) indicating that the sequenced isolate may be a polymorphic dikaryon or possibly a diploid. Repeat content in the two *Cryomyces* species varied from 8.4% in *C. minteri* to 12.5% in C. *antarcticus*, or about 2 Mbp, which also helps to explain the disparity in genome sizes between these species.

**Figure 1.**
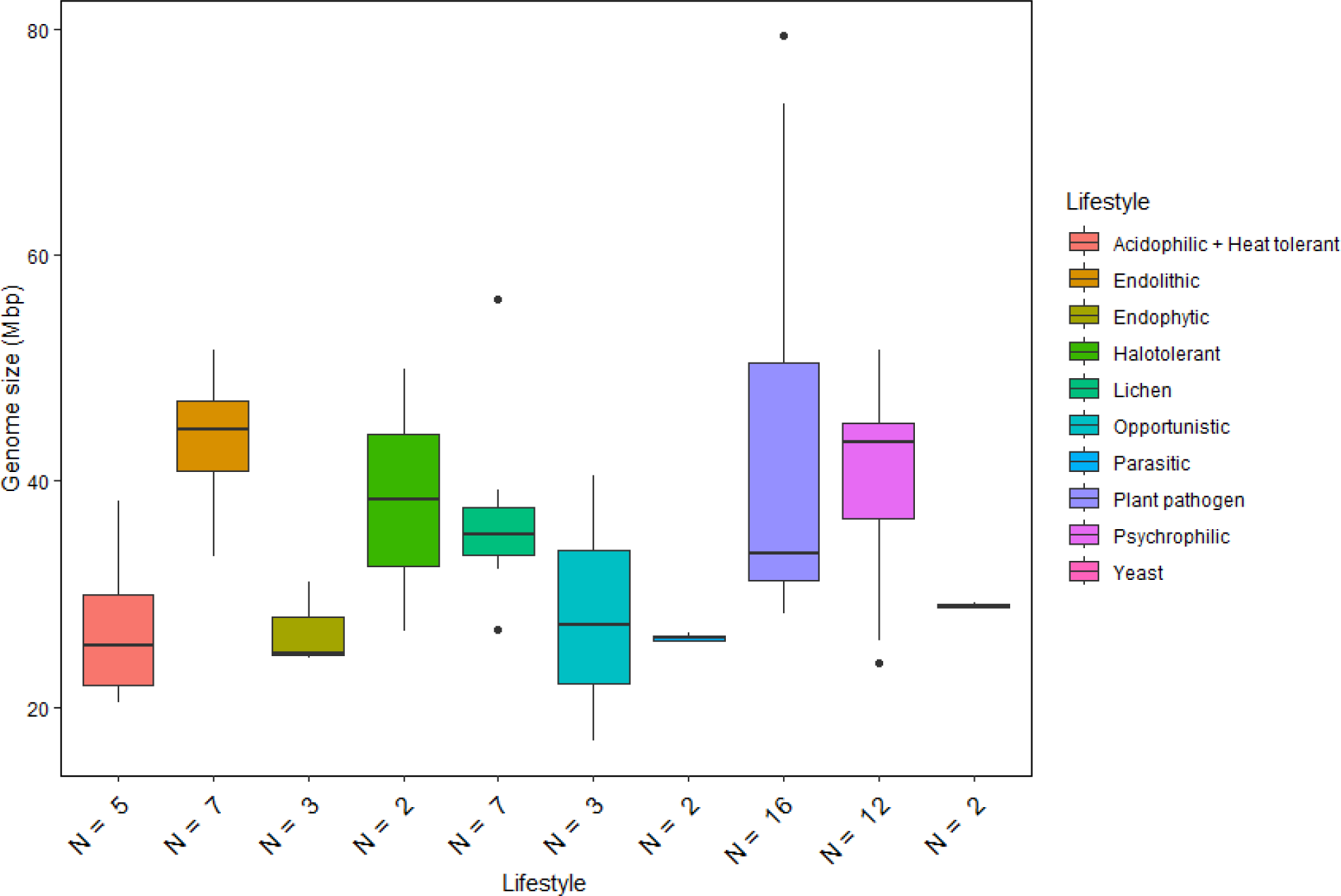
Genome size distribution of fungi with different lifestyles. Horizontal lines indicate the means, boxes show the group sizes, the vertical lines represent the range of the data, and the dots are outliers. Lifestyle groups are color coded according to the legend on the right.

**Figure 2.**
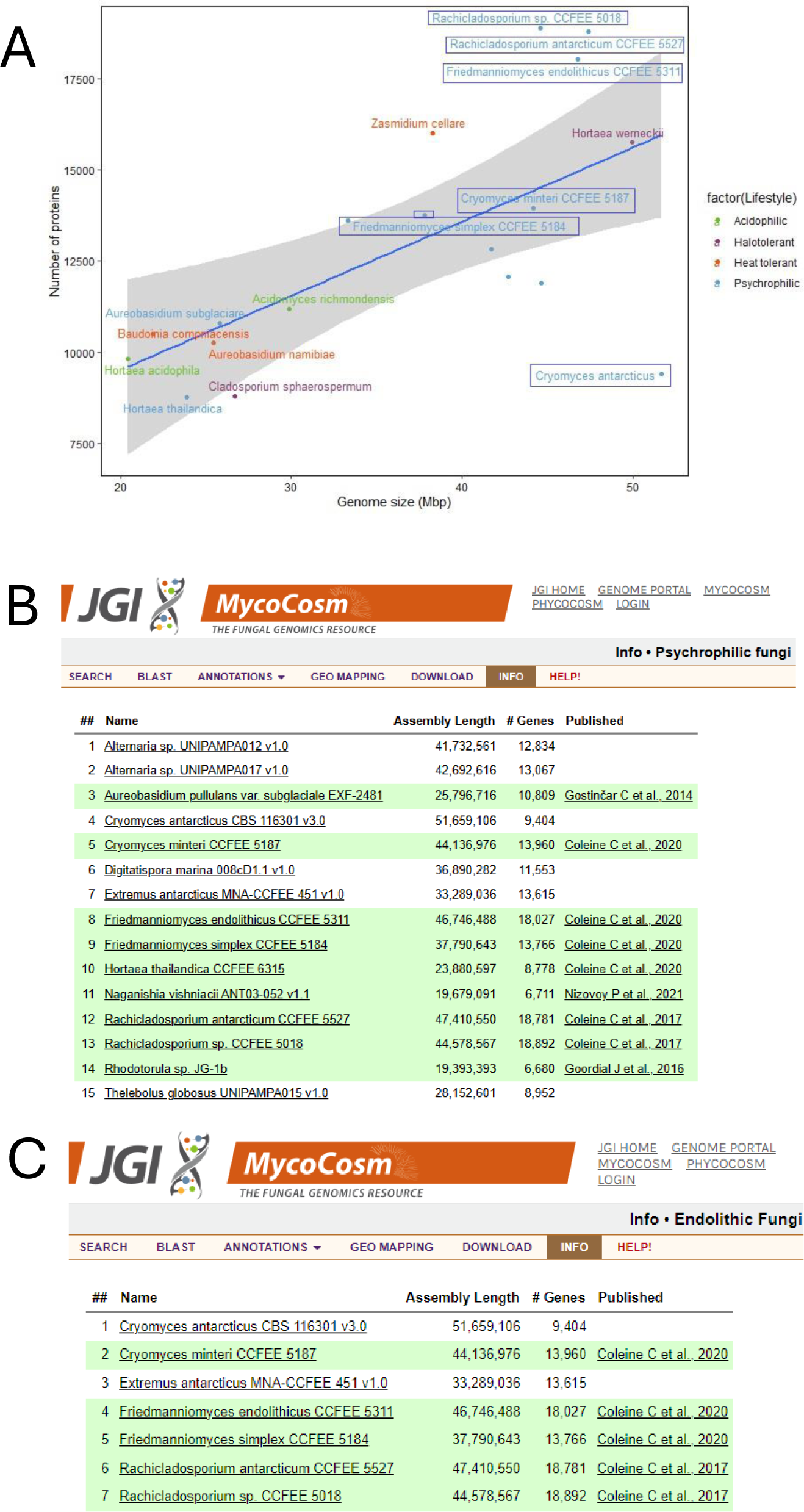
Summary information about the genomes of extremophilic fungi in MycoCosm, (**A**) Genome sizes compared to the numbers of encoded proteins for the extremophilic fungi. Each group is color coded according to the legend on the right. Species framed indicate the psychrophiles that are also endolithic. **(A-B)** MycoCosm group pages for for (**B**) psychrophilic (https://mycocosm.jgi.doe.gov/Psychrophilic_fungi) and (**C**) endolithic (https://mycocosm.jgi.doe.gov/Endolithic_fungi) groups of fungi with sequenced and annotated genomes showing genome assembly sizes, gene counts, and genome publication status.

**Figure 3.**
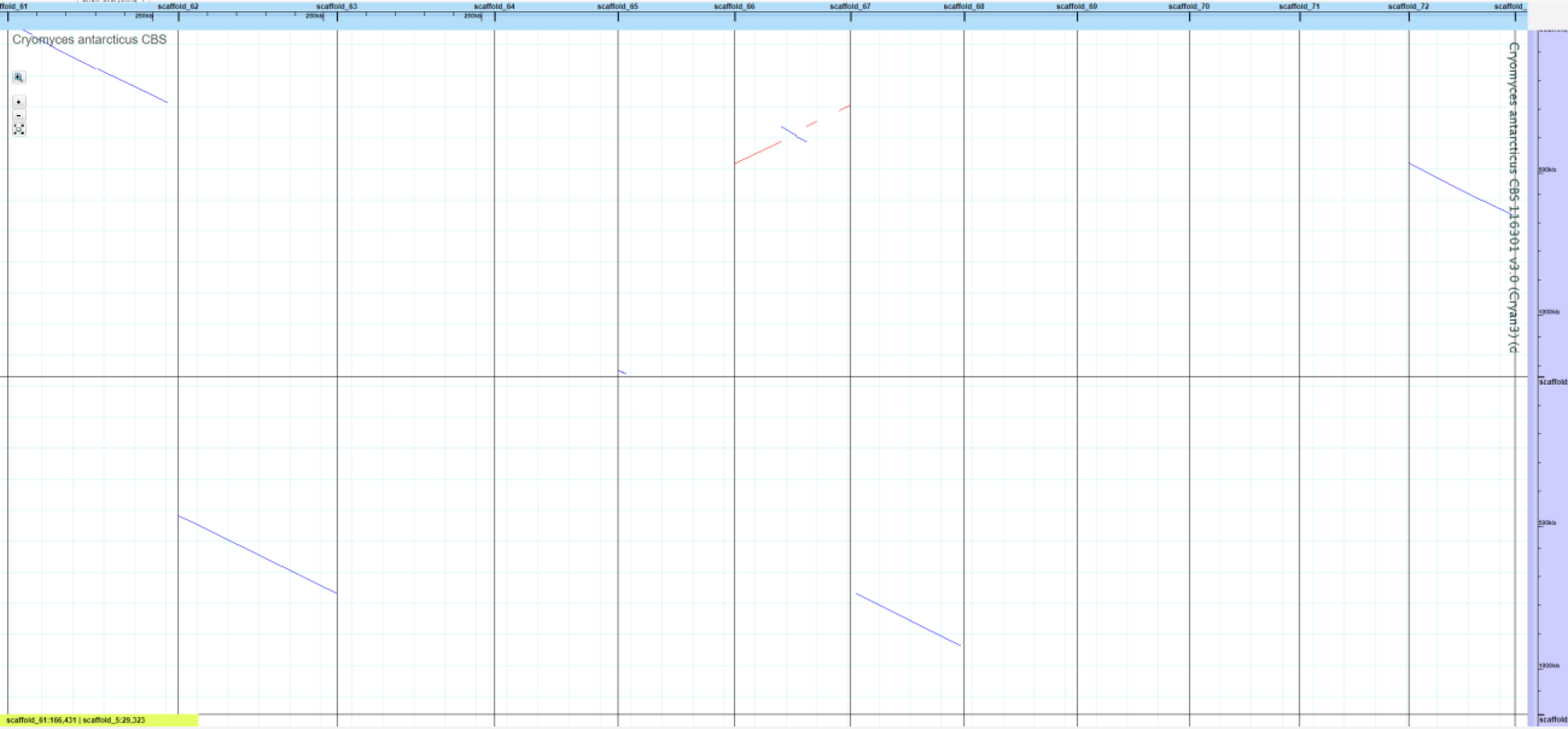
VISTA dot plot of selected *Cryomyces antarcticus* contigs from the whole-genome DNA-based self-alignment (Dubchak, 2007) – see Suppl Fig 1 for complete picture. The red and blue diagonal lines are nucleotide matches between different contigs, suggesting DNA duplication.

We also compared the genome sizes of the psychrophilic species with those of other lifestyle groups included in our analysis (Figure 1). The comparison revealed significant variations in genome size, with fungal pathogens generally exhibiting the largest genome sizes among the Dothideomycete species included in the study. Notably, the psychrophilic (including endolithic) and halotolerant fungi exhibit similar ranges of genome sizes, while other extremophiles classified as acidophilic and heat tolerant have reduced genome sizes compared to the other fungi.

There is a positive correlation between genome size and the number of encoded proteins among extremophilic fungi (Figure 2), where larger genome sizes corresponded to higher protein counts. However, there was a departure from this trend in six psychrophilic fungi, which deviated from the linear regression line. Notably, *F. endolithicus, Rachicladosporium* sp., and *R. antarcticum* show higher-than-expected protein counts relative to their genome sizes, which *C. antarcticus* show lower-than-expected protein counts. Despite this difference, we did not observe a consistent pattern genome/proteome ratio among all psychrophilic or extremophilic fungi when compared to other lifestyles. This suggests that the discrepancies in genome sizes compared to proteome size for some psychrophilic fungi may be unique to the individual species rather that indicative of general mechanisms of adaptation to extreme environments.

### 3.2 GC content in *Cryomyces* genomes

The *C. antarcticus* genome exhibited a bimodal distribution of GC content across contigs (Figure 4). The main peak of GC content contains a higher proportion of contigs with GC content values between 42% and 60%, while a small peak of low GC content between 0% and 32% contains a smaller number of contigs. This may be associated with some residual contamination or the presence of RIP mutations in repetitive regions, giving rise to low GC content. In *C. minteri,* we did not observe a clear bi-modal distribution of GC content as we did for *C. antarcticus*, potentially reflecting its lower proportion of repetitive DNA. We then analyzed the genome-wide RIP mutations in the genomes and found that both *C. minteri* and *C. antarcticus* showed a similar percentage of genome-wide RIP mutations of 4.36% and 4.04%, respectively, as calculated with the RIPper tool (Figure 5, Supplementary table 1). These values are the highest for total estimated genome-wide RIP among the psychrophilic species in this study. Taken together, these results suggest that the genomes of these two psychrophilic species might have been affected by RIP to some extent and at a higher level than other psychrophilic fungi. However, once again we did not observe a consistent pattern of genome-wide RIP or associated metrics across all psychrophilic fungal genomes. This suggests that this mechanism may not represent a general strategy for adaptation to cold stress.

**Figure 4.**
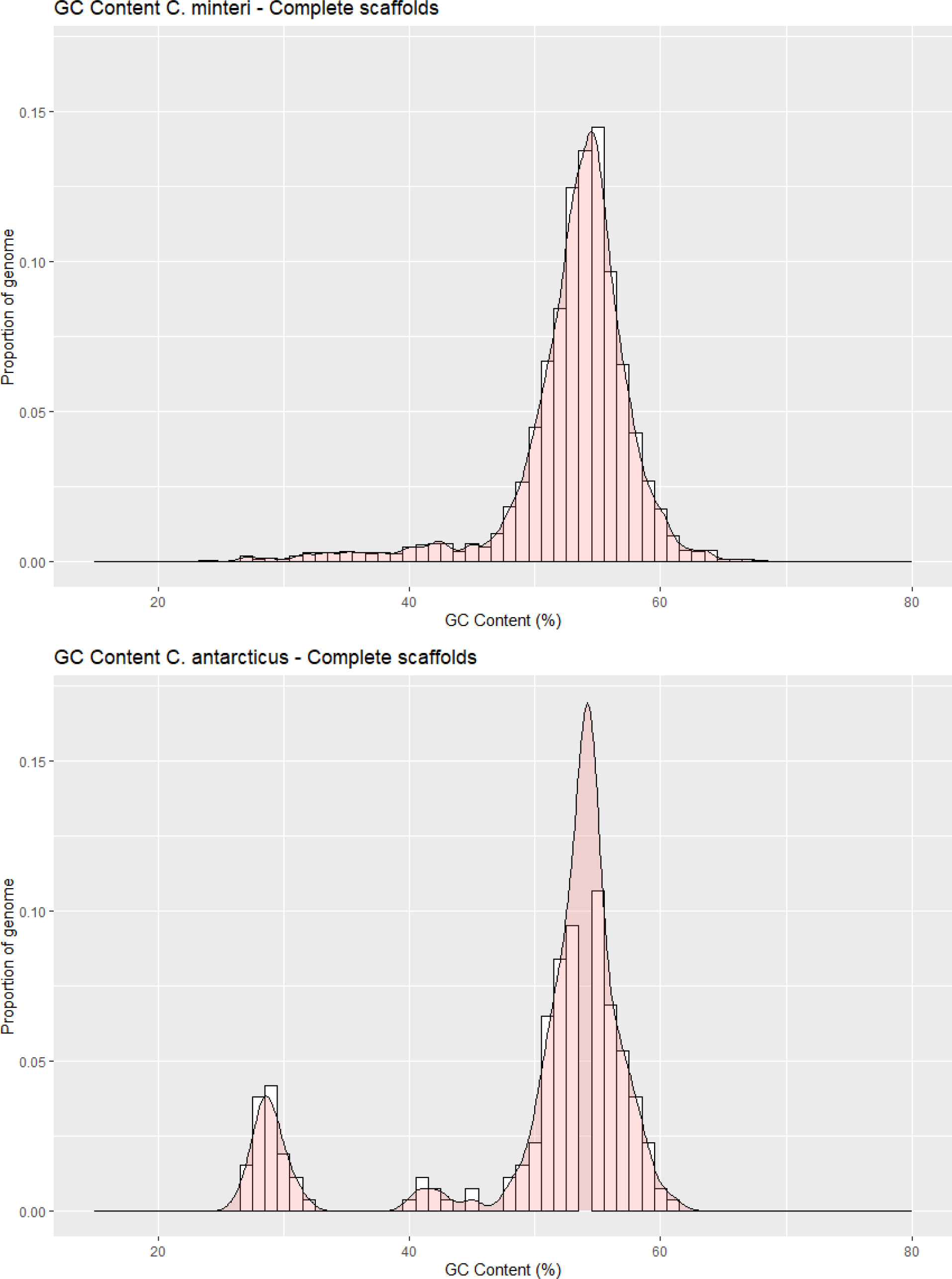
GC contents in the genomes of a) *Cryomyces minteri* and b) *C. antarcticus*

**Figure 5.**
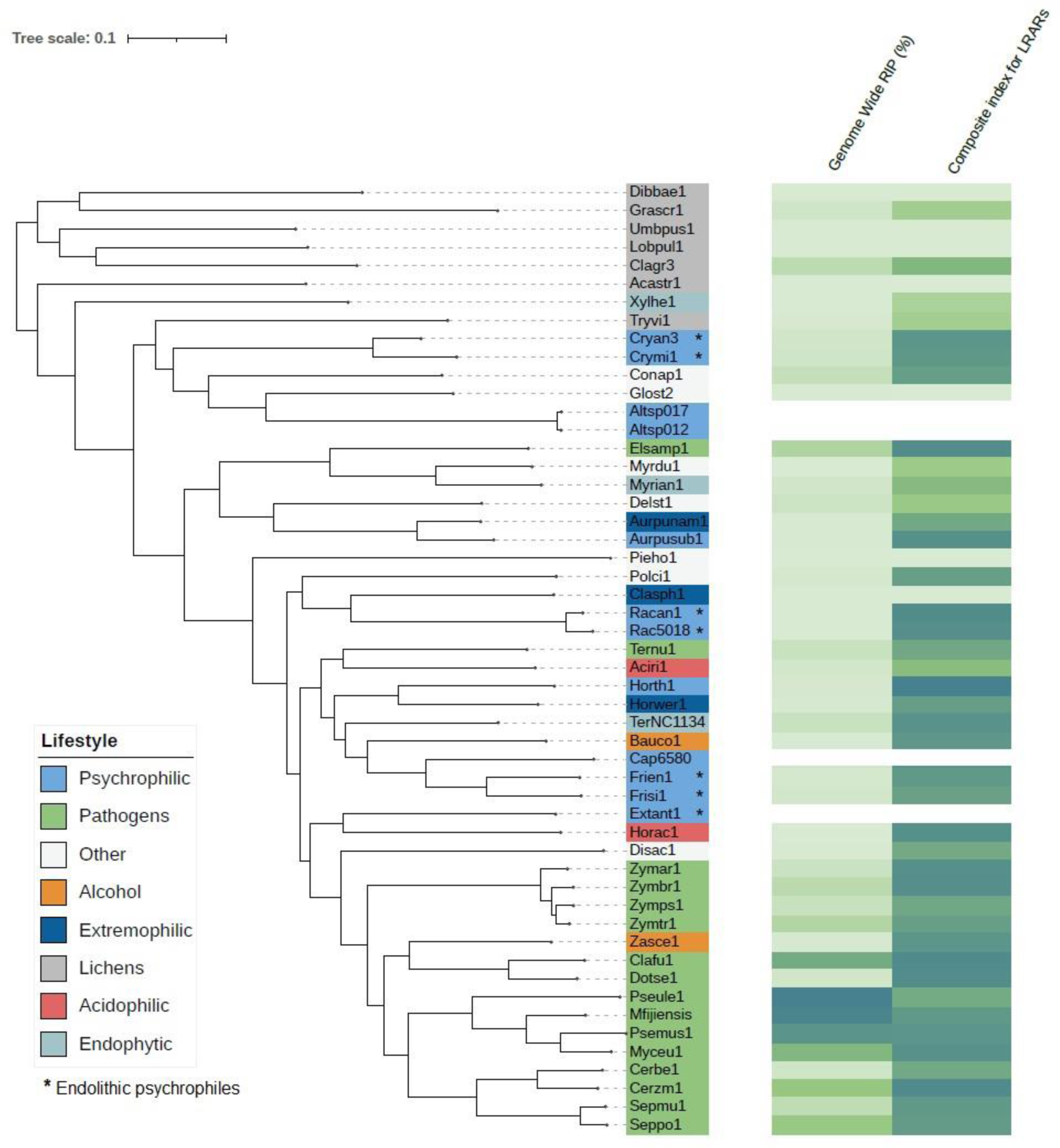
Phylogenetic tree produced by OrhthoFinder showing the percentage of genome-wide affected by RIP and the composite index for LRARs in each species. Tree leaves are labelled by JGI portal ids (see Table 1 for full species names).

### 3.3 GC content in whole-genome, protein-coding regions and GC3 is higher in psychrophilic and extremophilic fungi

The overall GC content in *C. antarcticus* and *C. minteri* is 53.31%. and 53.89%, respectively. While these values are consistent with those of fungi from other lifestyles, a broader analysis reveals some trends. Specifically, when considering the entire group of psychrophilic and extremophilic fungi, we observed that GC content in the psychrophiles (average = 54.38%) and extremophiles (average=53.47%) is higher than that in plant pathogens (average=50.38%), lichens (average = 47.74%) and fungi with other lifestyles (average = 51.02%). Further examination revealed that the GC content of coding sequences and GC content in the third-codon position (GC3) in the psychrophilic and extremophilic fungi are even higher than those in the fungi with other lifestyles (Figure 6), suggesting a pattern of elevated GC content in extremophiles across all DNA types.

**Figure 6.**
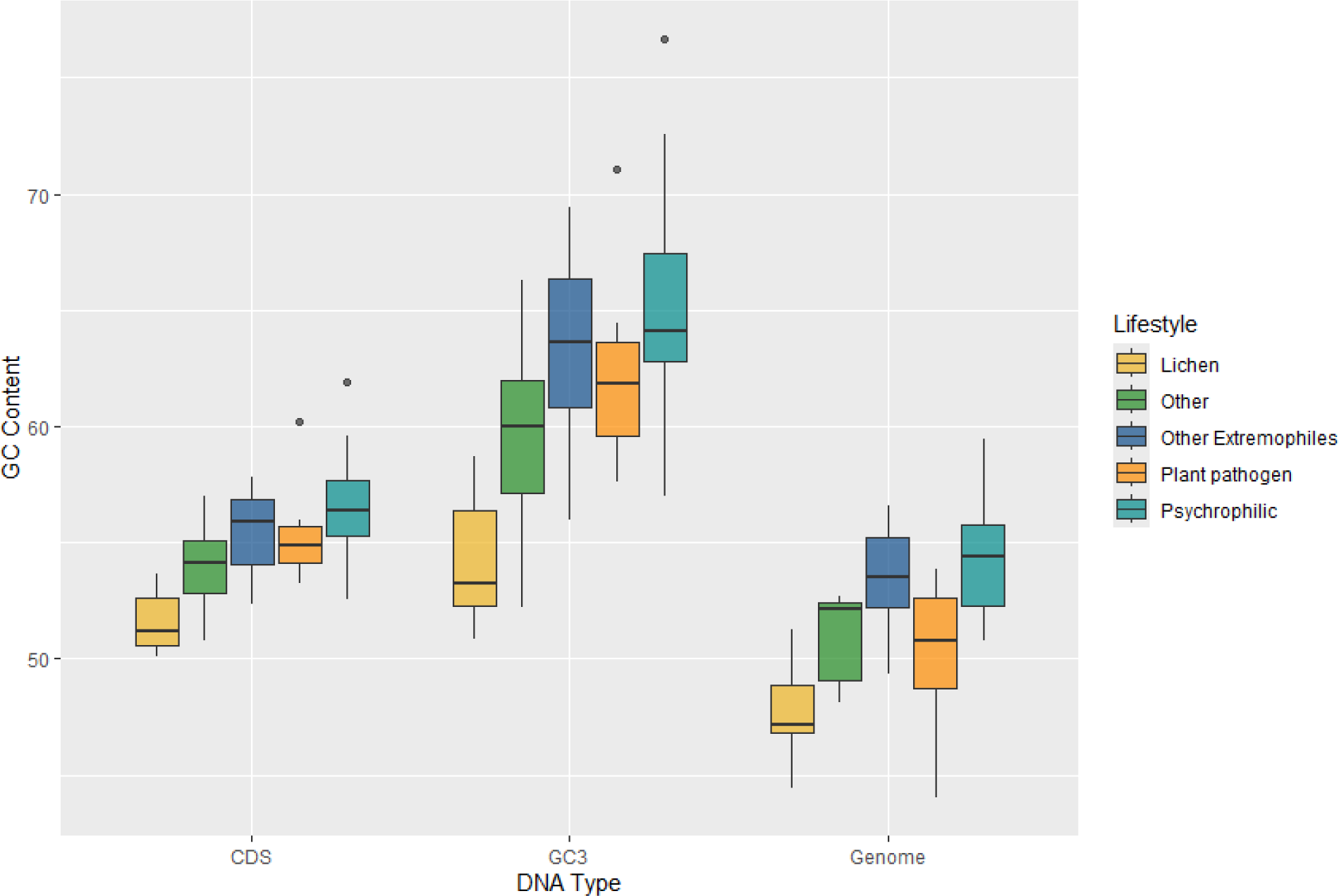
Plot of GC contents for coding sequences (CDS), third-position bases of genes (GC3) and the total genomes (Genome) for fungi in the different lifestyle groups, which are colored according to the legend on the right. Horizontal lines indicate the means, boxes show the group sizes, the vertical lines represent the range of the data, and the dots are outliers. Lifestyle groups are color coded according to the legend on the right.

### 3.4 OrthoFinder results

OrthoFinder assigned 616,448 genes from 52 species to 25,761 orthogroups. The number of orthogroups with all species present was 1,968, revealing the core genome of the fungal species. We identified 40 species-specific orthogroups for *C. antarcticus*, all of which have at least 2 paralogs in the species, and one orthogroup that has 4 members in *C. antarcticus*. In the case of *C. minteri,* we identified 148 species-specific genes, all of which are present in at least 2 copies, and in some cases even 4 to 8 copies. Overall, we found 233 genus-specific orthogroups in *Cryomyces*.

### 3.5 Expansion and contraction events in gene families distinguish psychrophilic and extremophilic fungi from other lifestyles

The total number of expanded and contracted orthogroups in the fungal species revealed more expansions and fewer contractions in gene families for psychrophilic fungi, while for fungi with other lifestyles, for instance some plant pathogens and lichens, the trend shows more contracted than expanded orthogroups in general (Figure 7). The heatmap clustering analysis reveals three distinct groups within the fungal species, which are due to the correlations between expanded and contracted orthogroups across all 52 species (Figure 8).

**Figure 7.**
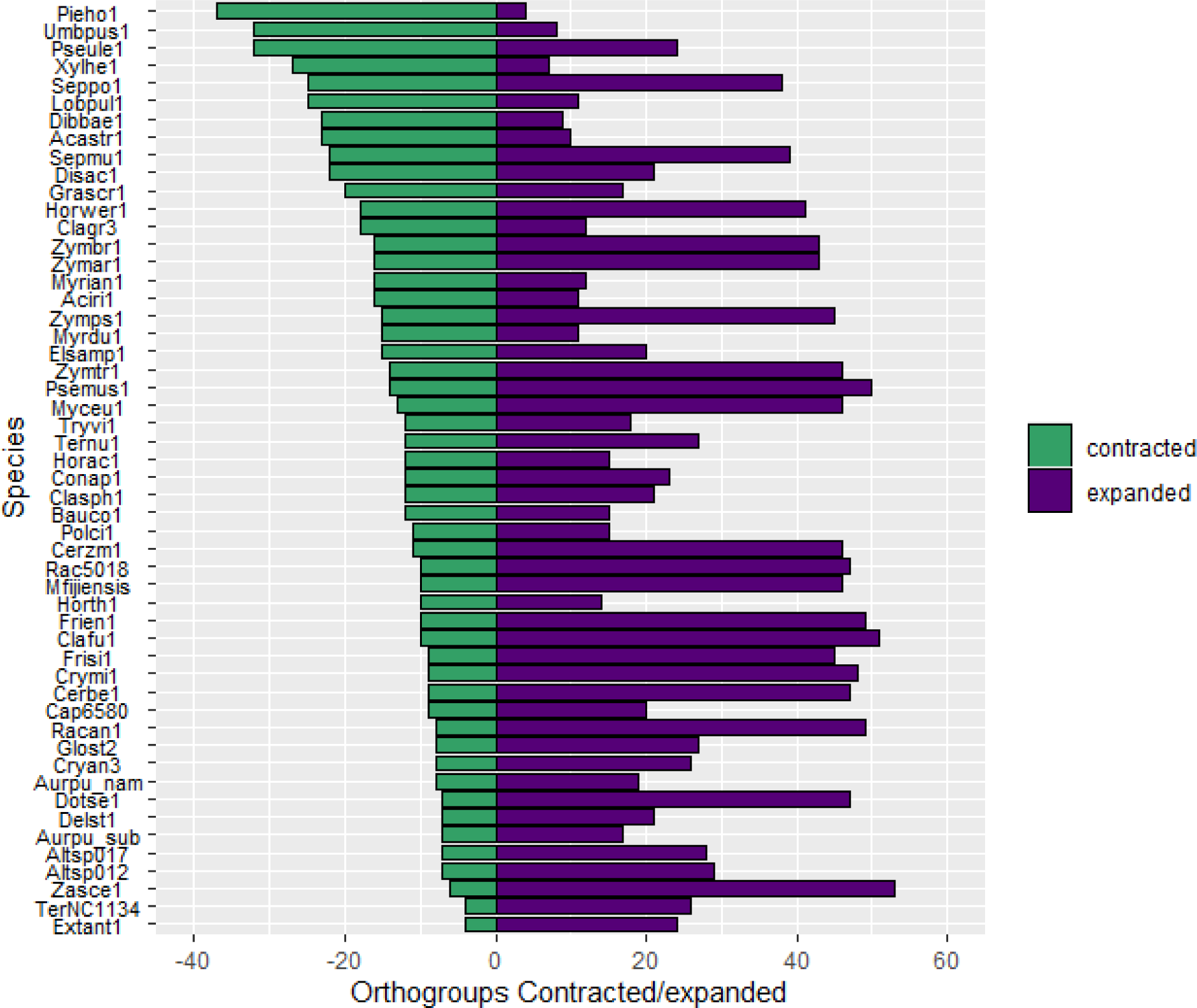
Numbers of orthogroups that are expanded (purple bars) or contracted (green bars) relative to a basal point of zero, no expanded/contracted. Groups are considered to be expanded or contracted in t > abs(0.75). Species are indicated by JGI portal ids (see Table 1 for full species names).

**Figure 8.**
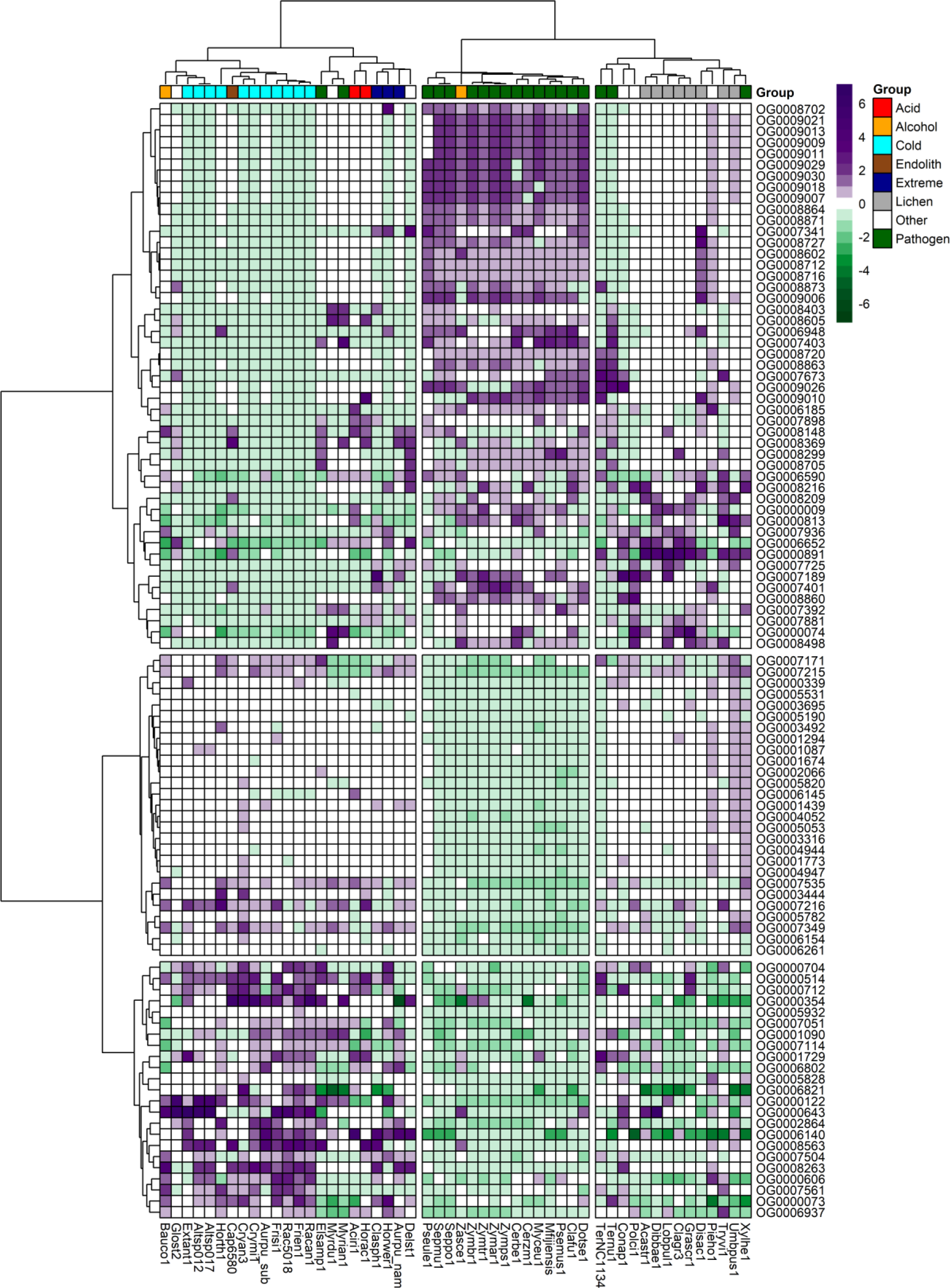
Heatmap of the top expanded and contracted orthogroups in the target psychrophilic species compared to the background fungal species.

The first group encompasses the psychrophilic species, which notably are clustered with other extremophilic fungi, including the acidophilic, halotolerant, heat tolerant, and the ethanol vapor-tolerant *B. compniacensis*. This implies the existence of a common survival strategy for withstanding environmental conditions associated with cold temperatures among psychrophilic species. These strategies parallel those employed by extremophilic fungi to overcome a diverse array of stresses associated with life in extreme conditions. The heatmap shows the 50 most contracted and expanded orthogroups based on t-score values. There are 30 orthogroups that are expanded in the psychrophilic species but generally contracted in the fungal plant pathogens and lichen species. On the other hand, 49 orthogroups are consistently contracted in the psychrophilic species and expanded in the plant pathogens; while 16 out of these 49 orthogroups are expanded in the third group which encompasses mostly the lichens, along with endophytic, opportunistic, yeasts and molds (Figure 8.).

### 3.6 Functional annotation of expanded gene families in psychrophilic species reveals diversification of mechanisms for cold tolerance

We categorize the expanded orthogroups into five general categories (Table 2) based on their predicted protein functions. We found CAZymes, including glycosyl hydrolases and transferases. Furthermore, enzymes with diverse functions underwent expansion, including carboxylases, peptidases, oxidoreductases, alcohol dehydrogenases, Phytanoyl-CoA dioxygenase, and hydrolases. Additionally, we found gene families that function as transporters including at least three orthogroups annotated as major facilitator super family (MFS) transporters. Some expanded orthogroups in the psychrophilic species have functional annotations related to DNA and RNA regulation, such as helicases, Zn(2)-C6 transcription factor, G-patch proteins, and Gfd2/YDR514C-like proteins. In addition, a fifth group of annotations consists of stress-related proteins, including cell wall integrity and stress response (WSC) family and aquaporins. Finally, a few orthogroups were uncharacterized or contain domains of unknown function.

**Table 2.** Functional annotations of the top 30 orthogroups that were expanded in psychrophilic species based on t-score analysis.

| **Orthogroup** | **Uniprot50 Blast Annotation** | **Functional group** |
| --- | --- | --- |
| **OG0000073** | Aquaporin-1 |  |
| **OG0000122** | Cell wall integrity and stress response component (WSC) domain | Stress-related |
| **OG0000354** | Gfd2/YDR514C-like C-terminal domain | DNA/RNA regulation |
| **OG0000514** | Enoyl reductase (ER) domain<br>Alcohol dehydrogenase GroES-like domain | PKS |
| **OG0000606** | Major facilitator superfamily (MFS) domain | Membrane transporter |
| **OG0000643** | Clr5 domain (Only rep sequence has this annotation, no cluster members support) |  |
| **OG0000704** | Isochorismatase-like domain |  |
| **OG0000712** | NAD(P)-binding protein<br>Cyclin-dependent kinases regulatory subunit |  |
| **OG0001090** | 3-phytase<br>Phosphoglycerate mutase-like protein |  |
| **OG0001729** | MFS-type transporter prx5<br>Major facilitator superfamily (MFS) profile domain | Membrane transporter |
| **OG0002864** | MFS general substrate transporter<br>Major facilitator superfamily (MFS) profile domain | Membrane transporter |
| **OG0003444** | Chromo domain |  |
| **OG0005782** | BTB domain |  |
| **OG0005828** | Zn(2)-C6 fungal-type domain | DNA/RNA regulation |
| **OG0005932** | Pyruvate carboxylase |  |
| **OG0006140** | AB hydrolase-1 domain |  |
| **OG0006802** | Glycoside hydrolase family 1 protein | CAZyme |
| **OG0006821** | Uncharacterized protein |  |
| **OG0006937** | EthD domain | Oxidoreductase |
| **OG0007051** | Phytanoyl-CoA dioxygenase family protein | Stress-related |
| **OG0007114** | Uncharacterized protein |  |
| **OG0007171** | MFS general substrate transporter | Membrane transporter |
| **OG0007215** | G-patch domain | DNA/RNA regulation |
| **OG0007216** | Dienelactone hydrolase domain |  |
| **OG0007349** | Oxidoreductase |  |
| **OG0007504** | Peptidase M20 dimerisation domain |  |
| **OG0007535** | Helicase-associated domain<br>Helicase-like protein | DNA/RNA regulation |
| **OG0007561** | Alpha-mannosyltransferase alg11p protein<br>(only some members of cluster) |  |
| **OG0008263** | Succinate dehydrogenase assembly factor 4,<br>mitochondrial | PKS |
| **OG0008563** | Zn(2)-C6 fungal-type domain | DNA/RNA regulation |

### 3.7 Gene families contracted in psychrophilic fungi are mostly required for interaction with plants and microorganisms

We conducted annotation analysis of the gene families that were contracted in psychrophilic fungi. Remarkably, the same orthogroups that were expanded in psychrophiles (section 3.6) were also often expanded in plant pathogens. Our analysis reveals a diverse array of annotations, including proteins involved in pathogenicity, stress response, effector production, cell wall integrity, and production of secondary metabolites (Table 3). Interestingly, the Ecp2 effector gene family, which is widely distributed among Dothideomycetes, is expanded in the plant pathogens but contracted in psychrophiles. Several orthogroups are associated with cell wall integrity, featuring annotations such as cell wall proteins, hydrophobic surface binding proteins, and acetyltransferases. In terms of stress response, we found gene families annotated as methyltransferases, and NACHT domain-containing proteins. Furthermore, multiple gene families associated with pathogenicity were identified, including F-box proteins, extracellular membrane proteins, oxidases, and enzymes for cell wall degradation and nutrient uptake. A remarkable finding was the expansion of the Zn(2)-C6 fungal-type domain, which can act as an aflatoxin regulatory protein according to the Interpro annotation.

**Table 3.**
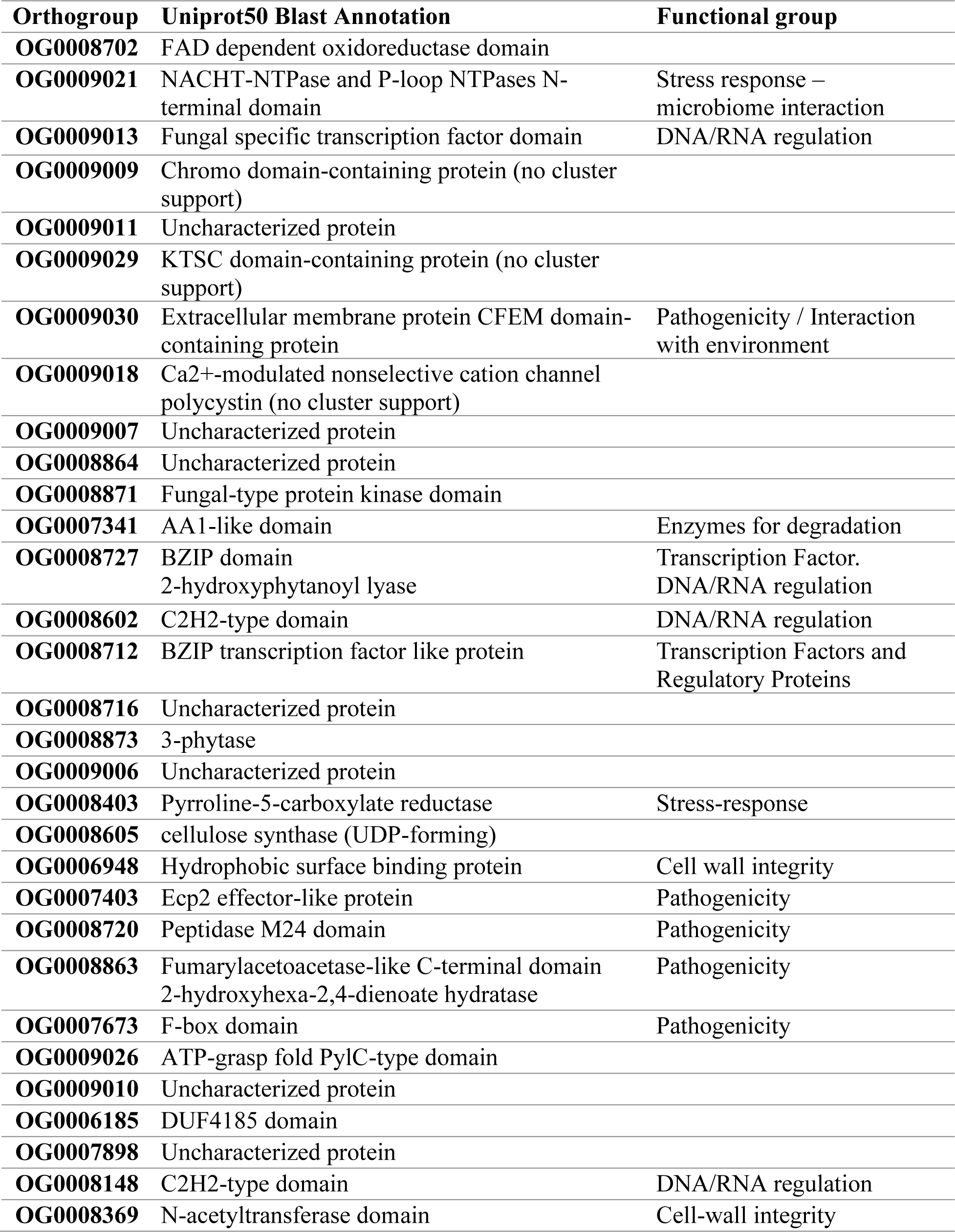
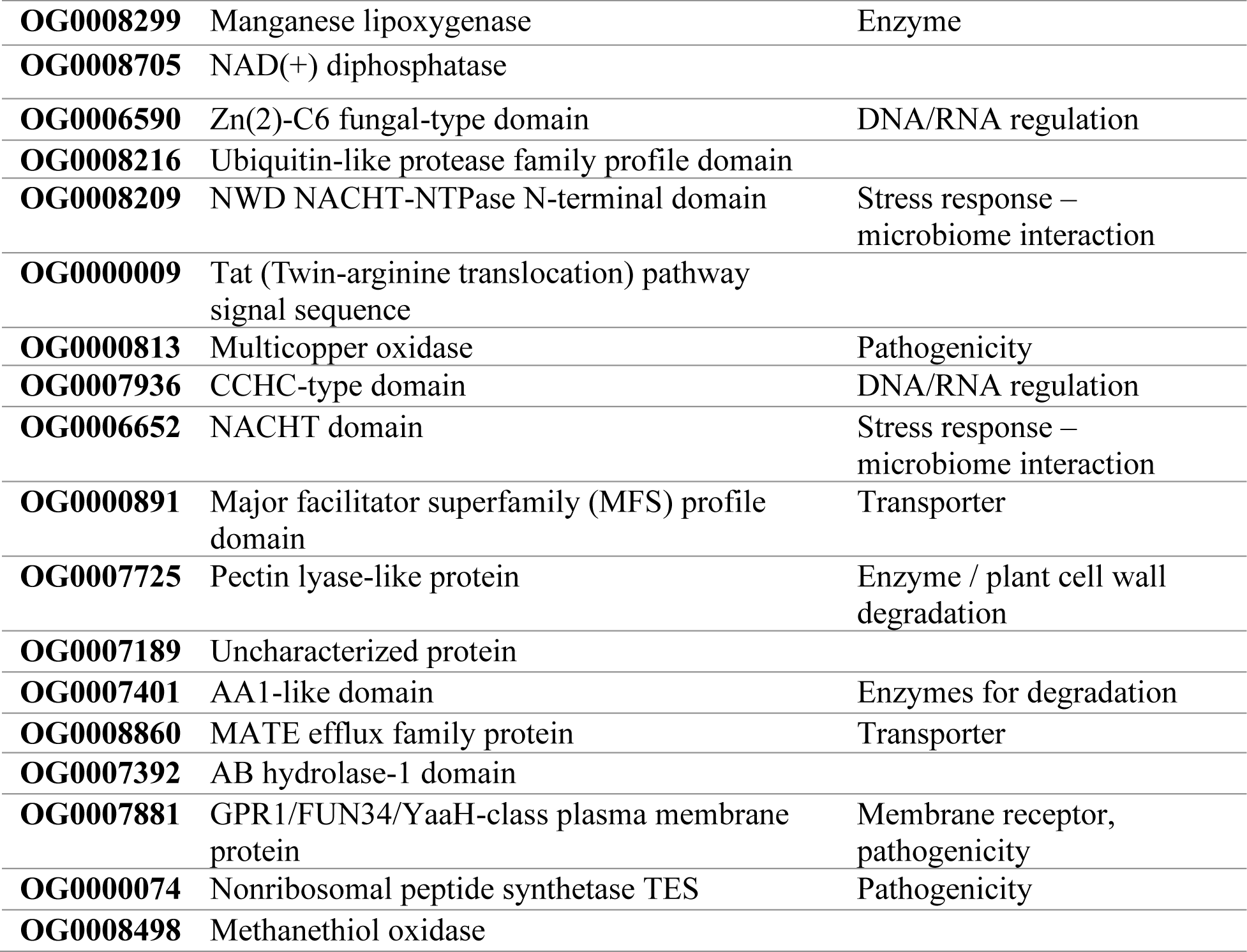
Functional annotations of the top 49 orthogroups that were contracted in psychrophilic species based on t-score analysis.

| **Orthogroup** | **Uniprot50 Blast Annotation** | **Functional group** |
| --- | --- | --- |
| **OG0008702** | FAD dependent oxidoreductase domain |  |
| **OG0009021** | NACHT-NTPase and P-loop NTPases N-terminal domain | Stress response – microbiome interaction |
| **OG0009013** | Fungal specific transcription factor domain | DNA/RNA regulation |
| **OG0009009** | Chromo domain-containing protein (no cluster support) |  |
| **OG0009011** | Uncharacterized protein |  |
| **OG0009029** | KTSC domain-containing protein (no cluster support) |  |
| **OG0009030** | Extracellular membrane protein CFEM domain-containing protein | Pathogenicity / Interaction with environment |
| **OG0009018** | Ca<sup>2+</sup>-modulated nonselective cation channel polycystin (no cluster support) |  |
| **OG0009007** | Uncharacterized protein |  |
| **OG0008864** | Uncharacterized protein |  |
| **OG0008871** | Fungal-type protein kinase domain |  |
| **OG0007341** | AA1-like domain | Enzymes for degradation |
| **OG0008727** | BZIP domain<br>2-hydroxyphytanoyl lyase | Transcription Factor.<br>DNA/RNA regulation |
| **OG0008602** | C2H2-type domain | DNA/RNA regulation |
| **OG0008712** | BZIP transcription factor like protein | Transcription Factors and Regulatory Proteins |
| **OG0008716** | Uncharacterized protein |  |
| **OG0008873** | 3-phytase |  |
| **OG0009006** | Uncharacterized protein |  |
| **OG0008403** | Pyrroline-5-carboxylate reductase | Stress-response |
| **OG0008605** | cellulose synthase (UDP-forming) |  |
| **OG0006948** | Hydrophobic surface binding protein | Cell wall integrity |
| **OG0007403** | Ecp2 effector-like protein | Pathogenicity |
| **OG0008720** | Peptidase M24 domain | Pathogenicity |
| **OG0008863** | Fumarylacetoacetase-like C-terminal domain<br>2-hydroxyhexa-2,4-dienoate hydratase | Pathogenicity |
| **OG0007673** | F-box domain | Pathogenicity |
| **OG0009026** | ATP-grasp fold PylC-type domain |  |
| **OG0009010** | Uncharacterized protein |  |
| **OG0006185** | DUF4185 domain |  |
| **OG0007898** | Uncharacterized protein |  |
| **OG0008148** | C2H2-type domain | DNA/RNA regulation |
| **OG0008369** | N-acetyltransferase domain | Cell-wall integrity |
| **OG0008299** | Manganese lipoxygenase | Enzyme |
| **OG0008705** | NAD(+) diphosphatase |  |
| **OG0006590** | Zn(2)-C6 fungal-type domain | DNA/RNA regulation |
| **OG0008216** | Ubiquitin-like protease family profile domain |  |
| **OG0008209** | NWD NACHT-NTPase N-terminal domain | Stress response –<br>microbiome interaction |
| **OG0000009** | Tat (Twin-arginine translocation) pathway<br>signal sequence |  |
| **OG0000813** | Multicopper oxidase | Pathogenicity |
| **OG0007936** | CCHC-type domain | DNA/RNA regulation |
| **OG0006652** | NACHT domain | Stress response –<br>microbiome interaction |
| **OG0000891** | Major facilitator superfamily (MFS) profile<br>domain | Transporter |
| **OG0007725** | Pectin lyase-like protein | Enzyme / plant cell wall<br>degradation |
| **OG0007189** | Uncharacterized protein |  |
| **OG0007401** | AA1-like domain | Enzymes for degradation |
| **OG0008860** | MATE efflux family protein | Transporter |
| **OG0007392** | AB hydrolase-1 domain |  |
| **OG0007881** | GPR1/FUN34/YaaH-class plasma membrane<br>protein | Membrane receptor,<br>pathogenicity |
| **OG0000074** | Nonribosomal peptide synthetase TES | Pathogenicity |
| **OG0008498** | Methanethiol oxidase |  |

## 4 Discussion

Previous analyses did not reveal much that was unique in the draft genome of a different isolate of *C. antarcticus,* and concluded with the question, “Nothing special in the specialist?” (Sterflinger et al., 2014). The previously identified haploid genome size and total GC contents were similar to our results. However, that genome was generated with short-read technology that yielded a very fragmented assembly containing over 12,400 contigs, compared to our PacBio assembly of isolate CBS 116301 of only 262 contigs. By greatly increasing the number of genomes from fungi with different lifestyles for comparative analyses it was possible to identify systematically expanded or contracted gene families in *C. antarcticus* and other psychrophiles that provide clues about their adaptations to extreme cold. Another result from our analysis is that many of the smaller contigs appear to represent an alternative haplotype to the major contigs, indicating that the sequenced isolate was likely diploid or dikaryotic. This is surprising because these fungi are believed to be haploid and asexual (Nieuwenhuis and James, 2016; Selbmann *et al*., 2020). Whether this represents a stage in the process of sexual reproduction, a balanced heterokaryon, or some other phenomenon is not known and should be the subject of future research, Due to differences in sequencing technologies caution should be applied when comparing contig-level genome assemblies, particularly when assessing features of genome evolution such as RIP, GC contents, and genome size. We observed a significant genome size difference between *C. minteri* and *C. antarcticus*, which is larger than expected given their close phylogenetic relationship but may be explained by differences sequencing and assembly methods, ploidy, and repeat content. A similar difference in genome size was observed in the genomes of the closely related pathogens *Epichloe typhina* and *E. clarkia,* where size discrepancies have been attributed to the impact of RIP on genome expansion (Treindl *et al*., 2021). Repeat-Induced Point mutation (RIP) functions as a defense mechanism in fungal genomes that counteracts transposable element activity and their proliferation (Gladyshev, 2017). This defense strategy induces C-to-T transition mutations within duplicated DNA sequences throughout the genome, resulting in the formation of GC-depleted regions and AT-rich sequences (Hood, Katawczik and Giraud, 2005; Testa, Oliver and Hane, 2016). RIP occurs during the sexual stage in haploid nuclei after fertilization but prior to meiotic DNA replication (de Wit *et al*., 2012). Our analysis revealed traces of RIP in the *C. minteri* and *C. antarcticus* genomes, covering roughly 4% of each genome. While the genome-wide RIP percentages observed in both *Cryomyces* species do not rank among the highest reported for fungi in the class Dothideomycetes, they do represent the highest within the psychrophilic fungi in our analysis and indicate that these species must undergo sexual reproduction at least occasionally.

Research on RIP and TE content in psychrophilic and extremophilic fungi is scarce. One study, focusing on the psychrophilic fungus *M. psychrophile,* revealed minimal repeat sequence content (Su *et al*., 2016). Currently, most of the research on RIP and TEs predominantly centers on plant pathogens. In fact, among the species included in our analysis, plant pathogens exhibited the highest RIP percentages, especially evident in the pathogens *P. ulei, M. fijiensis, P. musae, C. fulvum* and *M. eumusae,* which show more than 35% RIP. A high percentage of RIP has been reported previously in other fungal plant pathogens in the Dothideomycetes (de Wit *et al*., 2012; Ohm *et al*., 2012; Isaza *et al*., 2016). While research on extremophiles in this area is limited, the impact of RIP has been extensively explored in Dothideomycete plant pathogens including *Z. tritici* and its sister species, where a significant proportion of TEs exhibit RIP-like signatures (Lorrain *et al*., 2021). Analyses of *Z. tritici* isolates have revealed a higher incidence of RIP and TEs in accessory chromosomes compared to core chromosomes (Badet *et al*., 2020; Lorrain *et al*., 2021). Notably, *Z. tritici* maintains most of its effector genes on accessory chromosomes, often adjacent to TEs, indicating the crucial role of TEs in shaping the evolution of pathogenicity genes in fungi (Lorrain *et al*., 2021). Drawing parallels with plant pathogens, where key virulence genes are subject to RIP-induced changes, it is conceivable that RIP serves as a driver of gene evolution in extremophilic fungi, facilitating adaptation to extreme conditions. Given the necessity for environmental adaptation in extremophiles, similar to the importance of virulence in plant pathogens, RIP may contribute significantly to the evolutionary dynamics of genes involved in survival mechanisms.

GC content provides a simple overall indicator of nucleotide composition and can reveal variations in genomic stability and resistance to environmental stressors (Testa, Oliver and Hane, 2016; Hu *et al*., 2022). Studies have found that species with higher genome-wide GC content may exhibit greater resistance to high temperatures due to increased stability of their DNA helices (Hu *et al*., 2022; Liu *et al*., 2023). DNA stability is affected by temperature changes. There are two contributions to stability of DNA, enthalpy, and entropy. Entropy refers to single stranded polynucleotides that are coated with localized and partially localized water molecules and cations that are (at least partially) released when a double helix forms. There is a substantial increase in entropy (disorder) when the water and ions are released to the general solution. At low temperatures, water molecules are more organized, leading to more stability. Since the entropic destabilization decreases, adaptations to low temperature should prefer AT bases which have lower stability (enthalpy). This helps to maintain essential processes like replication and transcription to continue effectively. Contrary to the previous idea and to what we would expect, we observed higher GC content in genome, CDS and GC3 for psychrophiles. However, other studies have found similar results. For instance, in psychrophilic yeasts, the GC contents are higher than those of mesophilic yeasts (Liu *et al*., 2023). In the fungus *Mrakia psychrophila*, overall GC content surpassed that of mesophilic fungi such as *C. neoformans.* Also, the GC contents of the coding sequences (CDS) in two psychrophilic species, *M. psychrophile* and *D. cryoxerica,* were calculated at 56.5% and 56.8%, respectively. This reveals a higher percentage of GC in both psychrophiles than in *C. neoformans*, the reference mesophilic fungus in that study. A similar pattern, with even more pronounced differences, was observed for the third bases of codons (GC3) in both *M. psychrophile* and *D. cryoxerica*, compared to *C. neoformans* (Su *et al*., 2016). GC3 is also increased in thermophiles (Berka *et al.,* 2011). We hypothesize that the evidence suggesting a general trend of higher GC content in psychrophilic fungi, might be explained due to GC-rich regions having higher melting temperatures, which can protect the DNA from denaturation or damage caused by cold temperatures. This will suggest that while for some processes such as replication and transcription, a higher AT content is favorable, it is possible than a higher GC content helps to protect DNA from freezing damage caused by extreme low temperatures.

Expansions and contractions of gene families may provide additional clues about adaptations to an extreme environment. For example, we found expansions of helicases and G-patch domain-containing proteins, which are involved in multiple processes for genome regulation, in the psychrophilic species. In other organisms, DEAD-box RNA helicases are involved in regulation of nucleic acid binding, modulation of RNA/DNA:DNA interactions, and RNA metabolism, including translation initiation and ribosome biogenesis due to their capability to unwind the duplex RNA in eukaryotes (Lim, Thomas and Cavicchioli, 2000; Taschuk and Cherry, 2020). In psychrophilic microorganisms RNA helicases are differentially expressed at low temperature (Feller, 2013) and they can act as nucleic acid chaperones that help the microorganisms adapt to cold conditions (Xu, Xu and Liu, 2024). In particular, DEAD-box RNA helicases can presumably enable some bacteria to survive cold-shock and grow at low temperatures, due to their role in removing cold-stabilized secondary structures in mRNA (Lim, Thomas and Cavicchioli, 2000). In the Antarctic archaeon *M. burtonii,* an RNA helicase gene was highly transcribed during cell growth at 4 °C, revealing the importance of these proteins for low-temperature adaptation (Lim, Thomas and Cavicchioli, 2000). G-patch proteins are involved in regulation and activation of helicases. The first characterized protein with a G-patch domain that regulates an RNA helicase was Spp2 (Roy *et al*., 1995). The *in vivo* interaction of Spp2 was first documented in budding yeast, where a DEAH/RHA helicase named Prp2 interacts physically with Spp2, and this interaction is required for the activation of Prp2 function. Also, a helicase (Prp43) that is required in both splicing and ribosome biogenesis, is activated by a G-patch protein (Ntr1) which forms a stable complex with another G-patch protein (Ntr2). This complex recruits the helicase Prp43 targeting its helicase activity for spliceosome dissociation (Robert-Paganin, Réty and Leulliot, 2015).

Another gene family that was expanded in the psychrophiles was Zn(2)-C6 transcription factors, which are exclusively found in fungi. They can act as repressors, activators, or both, depending on the target genes (MacPherson, Larochelle and Turcotte, 2006). The regulatory scope of these transcription factors is broad, including activation of sterol biosynthesis (Vik and Rine, 2001), modulation of multidrug sensitivity (Coste *et al*., 2004), and regulation of genes involved in galactose metabolism (Breunig, 2000). Previous analyses have highlighted the versatility of Zn(2)-C6 transcription factors. It is particularly interesting that, in *Colletotrichum lagenarium*, Cmr1p positively regulates the expression of genes involved in melanin biosynthesis, which is crucial for successful invasion of host plants during anthracnose of cucumbers (Tsuji *et al*., 2000). In rock-inhabiting psychrophilic fungi, melanin biosynthesis is crucial for survival because melanin pigments protect the cells from extreme temperatures, solar irradiation, extreme osmotic and pH conditions, and provide tolerance to toxic metals (Selbmann *et al*., 2015). Moreover, Stb5, a Zn(2)-C6 transcription factor in *Saccharomyces cerevisiae*, can switch between activator and repressor roles under oxidative stress conditions (Larochelle *et al*., 2006). It binds to and regulates the expression of genes in the pentose phosphate pathway and those involved in NADPH production, which is essential for oxidative stress resistance. In fact, the activation of mechanisms for protection against oxidative stress plays a crucial role in psychrophilic fungi. It has been shown that a cold environment induces the production of reactive oxygen species (ROS), which can cause severe damage to DNA, lipids and proteins, affecting the cell survival (Kostadinova *et al*., 2017). The expansion of genes for Zn(2)-C6 transcription factors in *C. antarcticus* and other psychrophilic extremophiles may help to counter this damage.

Another expanded group includes major facilitator superfamily (MFS) transporters which likely are involved in drug resistance and amino acid uptake. The MFS are secondary transporters that have primary roles in antifungal resistance and nutrition transport. They can indirectly control membrane potential by changing membrane lipid homeostasis, and regulate internal pH and the stress response machinery in fungi (Su *et al*., 2016; Chen *et al*., 2017). MFS transporters are expanded in extremophilic fungal species such as *Mrakia psychrophila* and *Penicillium funiculosum* where they are required for accumulation of nutrients from the environment and resistance to acidic stress, respectively (Xu *et al*., 2014; Su *et al*., 2016). In *Alternaria alternata*, a MFS transporter named AaMFS19 is required for cellular resistance to oxidative stress, including H_2_O_2_, KO_2_, and other oxygen-generating compounds (Chen *et al*., 2017).

Some enzymes play a role in alternative pathways for energy production in fungi. We observed expansion of gene families of alcohol dehydrogenases. The expression of this family of enzymes has been reported, along with increased expression of glycolytic genes that feed into ethanol production via fermentation, in the Antarctic yeast *Rodothorula frigidialcoholis* growing at 0 °C (Touchette *et al*., 2022). This suggests that *R. frigidialcoholis* adapts to cold temperature by redirecting pentose phosphate pathway molecules to ethanol fermentation. This switch from respiration to fermentative metabolism to produce energy contributes to its survival in extreme cryoenvironments (Touchette *et al*., 2022). In *Mucor lusitanicus*, a knock-out strain of the alcohol dehydrogenase (*adh1*) gene, showed reduction in lipid and fatty acid content, which was restored by supplementing the media with external ethanol, suggesting that alcohol dehydrogenases are also involved in lipid biosynthesis (Gutiérrez-Corona *et al*., 2023). The expansion of genes for alcohol dehydrogenases in psychrophiles might be involved either in the activation of alternative pathways for energy production, or in specific stages of lipid biosynthesis, which both could be alternatives to live in extreme cold environments.

The expansion of CAZymes in the psychrophilic species may facilitate cell wall modifications and energy storage. Fungi living in cold environments maintain cell integrity by remodeling the cell wall. CAZymes control the repair and modification of cell wall components necessary to overcome extreme low temperatures and desiccation. For example, in *Candida albicans* the architecture of the cell wall undergoes alterations in response to environmental conditions, resulting in changes to its composition and structure in newly synthesized cells (Ene *et al*., 2015). In previous analyses, extremotolerant species have been shown to encode more members of the CAZyme family glycosyl hydrolase 32 (GH32) compared to opportunistic, non-opportunistic, and pathogenic species (Gostinčar, Zajc, Lenassi, Plemenitaš, de Hoog, *et al*., 2018). This family of CAZymes contains invertases and other enzymes involved in energy storage and recovery. In addition, studies have concluded that fungal cell wall elasticity is critical for survival under conditions of osmotic shock (Ene *et al*., 2015). Structural changes that improve cell wall elasticity involve cross-linking between cell wall macromolecules, which is catalyzed by the action of carbohydrate-active cell wall-remodeling enzymes, including hydrolases, transglycosidases, and transferases. These include the Gas-like family of β-1,3-glucanosyltransferases, and a family of chitin-glucanosyltransferases, which encompasses family GH16 (Ene *et al*., 2015; Ibe and Munro, 2021). In the fungal pathogen *B. cinerea*, the GT2 family contributes to the biogenesis and remodeling of the cell wall. This family contains the synthases for production of chitin, which plays a structural role in the fungal cell wall (Blandenet *et al*., 2022). In our analysis, we observe expansion of gene families of AB hydrolases (OG0006140), glycoside hydrolase family 1 (OG0006802), and the alpha-mannosyltransferase protein (OG0007561), which may be involved in cell wall modifications in response to cold temperatures.

Other expanded families, such as the phytanoyl-CoA dioxygenase proteins, are peroxisomal enzymes that catalyze the first step of phytanic acid alpha-oxidation. In *Streptomyces coelicolor,* an ectoine dioxygenase is involved in the synthesis of 5-hydroxyectoine, a compatible solute that is synthesized upon exposure to high salinity (Bursy *et al*., 2008). This compound helps organisms to survive extreme osmotic stress by acting as a highly soluble organic osmolyte.

Bursy *et al*. (2008) also found that synthesis of 5-hydroxyectoine by *S. coelicolor* is a stress response that protects against conditions of high salt and heat.

In contrast to gene families that were expanded in the psychrophiles, others were reduced, especially in comparison to the plant pathogens, in which gene families involved in microbiome interactions and defense or immune responses often were expanded. These families were contracted in psychrophilic fungi, possibly because plant pathogens require more proteins associated with microorganism and plant interactions, or interactions with other living things in general, while psychrophilic fungi are more isolated and face bigger challenges from abiotic and environmental factors rather than from other living organisms.

Most of the orthogroups that are contracted in the psychrophiles are proteins related to signaling and the membrane. They show diverse functions from degradation, response to stress, and synthesis of new compounds. Acetyltransferase (GNAT), heme haloperoxidase, and peroxidase are common domains in the contracted orthogroups. These proteins are involved in lignin and chitin degradation, which mostly are not used by cold-tolerant fungi (Leung, Robson and Robinson, 2011). However, the insect pathogen *Lecanicillium muscarium*, which was isolated from Antarctica, is a high producer of cold-tolerant enzymes, including those for hydrolyzing chitin, as expected for a fungus that feeds on arthropods (Fenice, 2016). Several fungi isolated from King George Island, Antarctica, including the Dothideomycetes *Penidiella kurandae*, showed amylase and cellulase activities, even when grown at 4 C (Krishnan *et al*., 2016).

Other reduced orthogroups in the psychrophilic fungi are associated with virulence/pathogenicity and toxin production in plant pathogens such as polyketide synthase, glycoside hydrolase, ribonuclease ribotoxin, aflatoxin regulatory protein, and pyrroline-5-carboxylate reductase. A previous comparative genomic analysis of 20 dothideomycetous and eurotiomycetous black fungi showed a lack of specialized virulence traits in extremotolerant and opportunistic fungi (Gostinčar, Zajc, Lenassi, Plemenitaš, De Hoog, *et al*., 2018). This is consistent with the contraction of these orthogroups that we observe.

We observed expansion of the NACHT-NTPase and P-loop NTPase N-terminal domain gene families in the group of plant pathogens, which suggests involvement of these proteins in controlling a variety of biotic interactions, not strictly limited to immune responses. The nucleotide-binding oligomerization domain (NOD)-like receptors (NLRs) are signal-transducing proteins that control innate immunity in plants, animals and fungi (Dyrka *et al*., 2014). In fungi, NACHT domain proteins, a subtype of NLRs, control immunity and other nonpathogenic biotic interactions, such as symbiotic relationships with microbiomes (Chu and Mazmanian, 2013). The HET-E protein of *Podospora anserina*, which first defined the NACHT domain, is involved in fungal non-self recognition and programmed cell death due to heterokaryon incompatibility (Saupe, Turcq and Bégueret, 1995; Koonin and Aravind, 2000). Additionally, studies have shown that NACHT domain proteins are specifically expressed during mycorrhizal symbiosis in certain fungi (Martin *et al*., 2008). These interactions could include fungal pathogenicity and symbiotic interactions (such as ECM formation, endophytic growth, lichen formation, or microbiome interactions).

In support of this idea, some studies suggest that variation in the number of NLR homologs among different fungal species might be associated with the diversity of ecological niches and lifestyles (Dyrka *et al*., 2014). For example, highly versatile pathogens like *Fusarium* species or mycoparasitic *Trichoderma* species tend to have large NLR repertoires, possibly due to protection against microbial competitors by the host immune system. However, the ethanol-tolerant *B. compniacensis* only has a small repertoire of these NLRs, which might result from inhabiting a restrictive niche (Dyrka *et al*., 2014). Similarly, our results suggest that extremophilic fungi which inhabit environments with a restricted number of microbial competitors and potential hosts, might not require specialized mechanisms for biotic interactions, which explains the contraction of the NACHT domain-containing protein family.

The contraction of hydrophobic surface-binding proteins among the psychrophiles may be due to the involvement of hydrophobins in water-sensing mechanisms during spore germination. Hydrophobins contributes to surface hydrophobicity needed for conidial dispersal, and protection against a host defense system, (Tanaka *et al*., 2022), all of which are essential processes for pathogenic or symbiotic interactions, but are not required in psychrophilic fungi. In plant pathogens like *M. grisea* and *F. graminearum,* hydrophobins are required for production of infective structures, penetration of the water-air interface of the host and attachment to hydrophobic surfaces (Talbot, Ebbole and Hamer, 1993; Quarantin *et al*., 2019). These properties are not required by saprobic, extremophilic psychrophiles and this is reflected in their proteome complements.

In addition to the gene families discussed above, we find that genes involved in stress responses against UV irradiation, oxidative stress, and antifungal agents, are also contracted, which suggests that the patterns of expansion and contraction do not necessarily favor a unique mechanism in each lifestyle. Instead, evolution and adaptation may have more to do with a dynamic and diversified set of strategies, as opposed to using only a specific set of proteins and mechanisms to cope with extreme environments.

While at first glance there was nothing special in the specialist (Sterflinger et al., 2014), upon a closer look with a larger cohort of fungi plus a much-improved genome sequence, expanded and contracted gene families can be identified and provide clues about adaptations to extreme conditions. This analysis highlights the importance of a large number of genomes available for comparison and validates the goals of projects like the JGI 1000 Fungal Genomes project (Grigoriev et al., 2014) which seeks to generate the requisite sequences. Clearly much additional work is needed to understand the expression of the identified gene families under extreme conditions to fully elucidate the ability of *Cryomyces* to grow under conditions that are more similar to those found on Mars than they are to Earth.

## Author contributions

SVG, MG and WS analyzed the data, prepared the figures and tables, and wrote the first draft of the manuscript. MG also helped select the genomes and guided the overall analysis. SH, KL, JE, NK, and AL contributed to sequencing, assembly, and annotation of the *Cryomyces* genome, coordinated by KB and IVG. IVG also initiated analysis and contributed to manuscript writing and editing. SBG spearheaded the generation of the *C. antarcticus* genome, provided project management, wrote several sections, and edited the entire manuscript. All authors have read and approved the final manuscript.

## Acknowledgements

We thank Dr. Filipe Victoria, Dr. Fabiola Lucini, and Dr. Francis Martin for providing access to the unpublished genome data of *Alternaria* sp. UNIPAMPA012 and *Alternaria* sp. UNIPAMPA017 produced within the 1000 Fungal Genomes project, the Dr. Laura Selbmann, the Italian National Program for Antarctic Researches (PNRA) and the Italian National Antarctic Museum (MNA) for providing access to the unpublished genome data of *Extremus antarcticus* MNA-CCFEE and *Capnodiales* sp, Dr. Elizabeth Arnold for providing access to the unpublished genome data of *Myriangiaceae* sp. NC1570 and *Teratosphaeriaceae* sp. NC1134, Dr. Ólafur Andrésson for providing access to the unpublished genome data of *Lobaria pulmonaria*, and Dr. Eduardo Mizubuti for providing access to the unpublished genome data of *Pseudocercospora ulei*. The genome sequence data were generated by the US Department of Energy Joint Genome Institute in collaboration with the user community. The work (proposal: 10.46936/10.25585/60001019) was conducted by the U.S. Department of Energy Joint Genome Institute (https://ror.org/04xm1d337), a DOE Office of Science User Facility, supported by the Office of Science of the U.S. Department of Energy operated under Contract No. DE-AC02-05CH11231. We also thank Jessica R. Cavaletto for the time-consuming process of extracting nucleic acids from the extremely slow growing and uncooperative *C. antarcticus*.

---

**Suppl. Figure 1.**
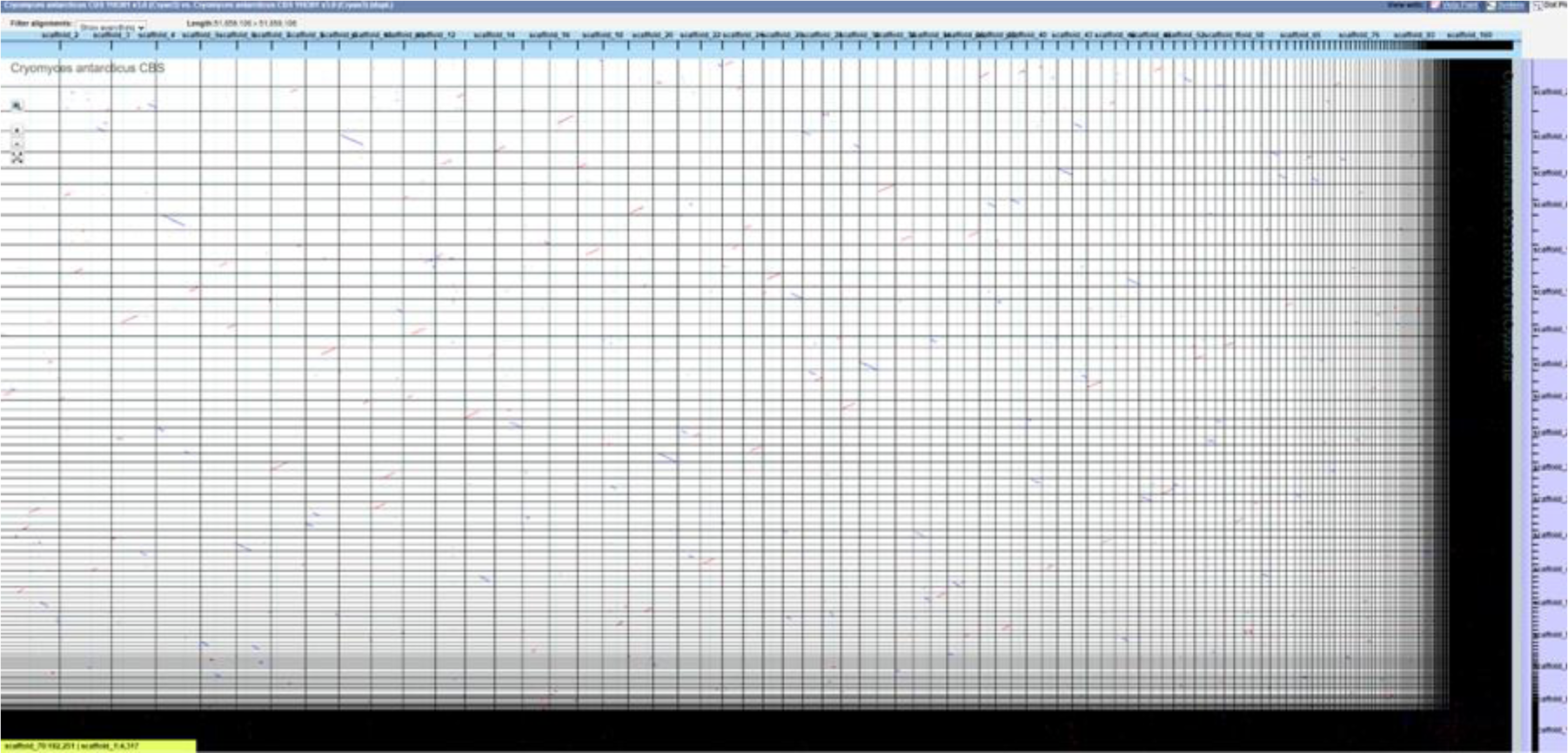
VISTA dot plot of *C. antarcticus* contigs against themselves based on the whole-genome DNA alignment (Dubchak, 2007). The red and blue diagonal lines are nucleotide matches between sequences on different contigs, indicating that the DNA duplication (the main diagonal of self-alignment has been removed for clarity).

**Supplementary table 1.** Repeat-induced point mutation (RIP) in the genomes used for comparative analysis.

| Species name | JGI<br>Portal<br>ID | Total<br>estimated<br>genome wide<br>RIP (%) | Average<br>size of<br>LRAR<br>(kbp) | Average<br>GC<br>content of<br>LRAR (%) | Sum of<br>all<br>LRAR<br>(Mbp) | Composit<br>e index<br>for<br>LRARs |
| --- | --- | --- | --- | --- | --- | --- |
| *Acarospora strigata* | Acastr1 | 0.09 | 0 | 0 | 0 | 0 |
| *Acidomyces richmondensis*<br>*BFW* | Aciri1 | 3.2 | 6.1 | 40.4 | 0.2 | 1.05 |
| *Aureobasidium namibiae* | Aurp_n<br>am1 | 0.77 | 7.38 | 37.27 | 0.1 | 1.32 |
| *Aureobasidium subglaciare* | Aurpu_<br>sub1 | 1.18 | 9.17 | 37.41 | 0.1 | 1.66 |
| *Baudoinia compniacensis* | Bauco1 | 0.78 | 8.98 | 32.43 | 0.1 | 1.57 |
| *Cercospora berteroa* | Cerbe1 | 4.75 | 7.27 | 40.6 | 0.2 | 1.32 |
| *Cercospora zae-maydis* | Cerzm1 | 29.67 | 13.32 | 36.75 | 9.2 | 1.73 |
| *Cladosporium fulvum* | Clafu1 | 42.2 | 10.07 | 42.18 | 10.3 | 1.73 |
| *Cladonia grayi* | Clagr3 | 12.61 | 12.72 | 15.8 | 2.4 | 1.11 |
| *Cladosporium sphaerospermum* UM 843 | Clasph1 | 0 | 0 | 0 | 0 | 0 |
| *Coniosporium apollinis* | Conap1 | 9.65 | 14.96 | 30.04 | 1.9 | 1.45 |
| *Cryomyces antarcticus* | Cryan3 | 4.04 | 10.97 | 39.27 | 1.5 | 1.58 |
| *Cryomyces minteri*<br>*CCFEE 5187* | Crymi1 | 4.36 | 9.42 | 37.36 | 1 | 1.55 |
| *Delphinella strobiligena* | Delst1 | 5.56 | 5.83 | 23.08 | 0.9 | 0.85 |
| *Dibaeis baeomyces* | Dibbae<br>1 | 0.21 | 0 | 0 | 0 | 0 |
| *Dissoconium aciculare* | Disac1 | 0.75 | 8.58 | 26.04 | 0.1 | 1.3 |
| *Dothistroma septosporum* | Dotse1 | 3.66 | 30.74 | 31.59 | 1.1 | 1.7 |
| *Elsinoë ampelina* | Elsamp<br>1 | 17.82 | 18.16 | 31.11 | 4.3 | 1.69 |
| *Friedmanniomyces endolithicus*<br>CCFEE 5311 | Frien1 | 2.4 | 9.04 | 42.57 | 0.7 | 1.52 |
| *Friedmanniomyces simplex*<br>CCFEE 5184 | Frisi1 | 3.62 | 7.15 | 41.68 | 0.3 | 1.43 |
| *Glonium stellatum* | Glost2 | 0.88 | 0 | 0 | 0 | 0 |
| *Graphis scripta* | Grascr1 | 4.31 | 5.94 | 23.55 | 0.4 | 0.74 |
| *Hortaea acidophila* | Horac1 | 0.29 | 4.5 | 32.87 | 0 | 1.66 |
| *Hortaea thailandica* | Horth1 | 1.84 | 9.8 | 46.55 | 0.2 | 1.85 |
| *Hortaea werneckii* | Horwer1 | 1.32 | 7.85 | 37.07 | 0.4 | 1.47 |
| *Lobaria pulmonaria* | Lobpul1 | 0.06 | 0 | 0 | 0 | 0 |
| *Mycosphaerella fijiensis* | Mfijien<br>sis_v2 | 59.25 | 29.42 | 40.44 | 40.7 | 1.52 |
| *Mycosphaerella eumusae* | Myceu1 | 36.49 | 9.57 | 40.9 | 4.4 | 1.64 |
| *Myriangium duriae* | Myrdu1 | 0.23 | 5.75 | 20.42 | 0 | 0.81 |
| *Myriangiaceae*<br>sp. NC1570 | Myrian1 | 3.9 | 13.41 | 24.85 | 0.8 | 1.08 |
| *Piedraia hortae* CBS<br>480.64 v1.1 | Pieho1_1 | 0.02 | 0 | 0 | 0 | 0 |
| *Polychaeton citri* | Polci1 | 2.32 | 7.31 | 33.77 | 0.1 | 1.46 |
| *Pseudocercospora musae* | Psemus1 | 52.5 | 11.13 | 36.2 | 8.4 | 1.59 |
| *Pseudocercospora ulei* | Pseule1 | 60.76 | 9.65 | 47.32 | 36.4 | 1.28 |
| *Rachicladosporium* sp.<br>CCFEE 5018 | Rac5018_1 | 0.4 | 6.33 | 40.2 | 0 | 1.67 |
| *Rachicladosporium antarcticum*<br>CCFEE 5527 | Racan1 | 0.6 | 7.13 | 43.46 | 0.1 | 1.72 |
| *Septoria musiva* | Sepmu1 | 11.31 | 15.04 | 41.73 | 2.8 | 1.53 |
| *Septoria populicola* | Seppo1 | 28.45 | 12.71 | 43.84 | 4.7 | 1.51 |
| *Teratosphaeria*<br>*aceae* sp.<br>*NC1134* | TerNC1<br>134_1 | 7.84 | 14.24 | 31.67 | 1.1 | 1.61 |
| *Teratosphaeria*<br>*nubilosa* | Ternu1 | 7.49 | 9.15 | 27.07 | 1.1 | 1.35 |
| *Viridothelium*<br>*virens* | Tryvi1 | 1.52 | 7.17 | 16.61 | 0 | 0.71 |
| *Lasallia*<br>*pustulata*<br>(*Umnilicaria*<br>*pustulata*) | Umbpu<br>s1 | 0.43 | 0 | 0 | 0 | 0 |
| *Xylona heveae* | Xylhe1 | 0.38 | 4.83 | 17.14 | 0 | 0.6 |
| *Zasmidium*<br>*cellare* | Zasce1 | 1.76 | 6.44 | 37.2 | 0.1 | 1.58 |
| *Zymoseptoria*<br>*ardabilia* | Zymar1 | 6.32 | 7.45 | 41.2 | 0.3 | 1.66 |
| *Zymoseptoria*<br>*brevis* | Zymbr1 | 13.97 | 7.48 | 40.56 | 0.3 | 1.68 |
| *Zymoseptoria*<br>*pseudotritici* | Zymps1 | 8.46 | 8.26 | 43.46 | 0.5 | 1.35 |
| *Zymoseptoria*<br>*tritici* | Zymtr1 | 17.62 | 11.17 | 43.43 | 6.2 | 1.46 |

